# Cofilin regulates actin network homeostasis and microvilli length in mouse oocytes

**DOI:** 10.1101/2021.10.15.464498

**Authors:** Anne Bourdais, Benoit Dehapiot, Guillaume Halet

## Abstract

How multiple actin networks coexist in a common cytoplasm, while competing for a shared pool of monomers, is still an ongoing question. This is exemplified by meiotic maturation in the mouse oocyte, which relies on the dynamic remodeling of distinct cortical and cytoplasmic F-actin networks. Here we show that the conserved actin-depolymerizing factor cofilin is activated in a switch-like manner at meiosis resumption from prophase arrest. Interfering with cofilin activation during maturation resulted in widespread microvilli elongation, while cytoplasmic F-actin was depleted, leading to defects in spindle migration and polar body extrusion. In contrast, cofilin inactivation in metaphase II-arrested oocytes resulted in a shutdown of F-actin dynamics, along with a dramatic overgrowth of the polarized actin cap. However, inhibition of the Arp2/3 complex to promote actin cap disassembly elicited ectopic microvilli outgrowth in the polarized cortex. These data establish cofilin as a key player in actin network homeostasis in oocytes, and reveal that microvilli can act as a sink for monomers upon disassembly of a competing network.

## Introduction

Actin filaments (F-actin) control a vast array of cellular processes requiring membrane deformation or force production, such as cell motility and contractility, cell division, vesicular trafficking and cell polarization (Pollard and Cooper, 2009). Multiple actin networks coexist in the cell, exhibiting elaborate architectures that are defined primarily by their nucleation machinery (Michelot and Drubin, 2011; Blanchoin et al., 2014). Thus, the Arp2/3 complex generates branched actin networks by promoting the nucleation of new filaments from the side of existing ones (Mullins et al., 1998). In contrast, formins nucleate and/or elongate linear actin cables, to promote the formation of e.g. stress fibers, filopodia and the cytokinetic ring (Chesarone et al., 2010). An emerging concept in the field is internetwork competition, by which distinct actin structures compete for a finite pool of actin monomers (G-actin), thereby regulating network size and density (Burke et al., 2014; Suarez and Kovar, 2016). How this homeostatic balance is regulated in living cells is not fully understood. So far, emphasis has been placed on the relative amounts, and the assembly rates, of each family of nucleators (i.e. Arp2/3 vs formins), along with the key role of profilin in allocating G-actin toward formins at the expense of Arp2/3 (Rotty et al., 2015; Suarez et al., 2015; Antkowiak et al., 2019; Faust et al., 2019). However, little is known of the contribution of F-actin disassembly mechanisms, particularly in a cellular context.

Actin filaments are structurally polarized, with a fast growing barbed end and a slow growing pointed end (Blanchoin et al., 2014). In vivo, the barbed end is the preferred site for monomer addition and filament elongation, while the pointed end favors monomer disassembly, resulting in a directional shuttling of actin subunits known as treadmilling (Carlier and Shekhar, 2017). Actin-binding proteins of the ADF/cofilin family (herein, cofilin) play an instrumental role in actin network turnover, by promoting filament severing and depolymerization from the pointed end, thereby replenishing the cytoplasmic pool of monomers (Lappalainen and Drubin, 1997; Hotulainen et al., 2005; Suarez et al., 2011; Shekhar and Carlier, 2017; Wioland et al., 2017). In addition, cofilin lowers the affinity of the Arp2/3 complex for both the mother and daughter filaments, resulting in debranching and depolymerization of dendritic networks (Blanchoin et al., 2000; Chan et al., 2009). On the other hand, if not rapidly capped, the free barbed ends generated through severing could seed new filament growth (Chan et al., 2000; Ichetovkin et al., 2002). Thus, cofilin plays a dual role in actin homeostasis, promoting either a net increase, or decrease, in F-actin density. Phosphorylation at Ser3, catalyzed by LIM kinases (LIMK), prevents cofilin binding to actin, and thereby inhibits its filament depolymerizing activity (Arber et al. 1998; Yang et al., 1998; Mizuno, 2013).

Oocytes offer an excellent experimental paradigm to address actin network homeostasis at the cell scale. During meiotic maturation, mouse oocytes experience a profound remodeling of their actin cytoskeleton to achieve symmetry breaking and polar body extrusion (reviewed in Uraji et al., 2018; Duan and Sun, 2019). Eccentric positioning of the first meiotic spindle relies on a dynamic network of cytoplasmic actin filaments nucleated by Fmn2 (Azoury et al., 2008; Schuh and Ellenberg, 2008; Pfender et al., 2011; Yi et al., 2013). Intriguingly, the density of this actin mesh drops steeply at meiosis resumption, and its gradual reassembly drives spindle migration toward the oocyte cortex (Azoury et al., 2011; Holubcová et al., 2013; Wei et al., 2018). Fmn2-dependent actin filaments are also found along spindle microtubules (spindle actin), where they promote kinetochore fibre formation and chromosome alignment (Mogessie and Schuh, 2017). Spindle migration also relies on an Arp2/3-dependent cortical actin thickening, which contributes to decreasing cortical tension (Chaigne et al., 2013). In mature oocytes arrested in metaphase-II (MII), a prominent actin cap is established in the cortex overlying the spindle, thus defining the site for second polar body protrusion after fertilization (Longo and Chen, 1985). Filaments in the cap are assumed to be arranged as a branched network, as they rely on the polarized activation of the Arp2/3 complex (Yi et al., 2011). Remarkably, actin cap formation is invariably associated with the collapse of microvilli in the polarized cortex, but the molecular underpinnings of this cytoskeletal remodeling are still poorly understood (Maro et al., 1986; van Blerkom and Bell, 1986; Dehapiot and Halet, 2013).

In this study, we explored the role of cofilin in regulating actin network dynamics during mouse oocyte maturation. Unexpectedly, we found that cofilin is largely inactivated (ie, phosphorylated) during prophase arrest, and becomes active at meiosis resumption. Inhibition of cofilin during meiosis-I via LIMK1 overexpression resulted in a striking elongation of microvilli and a loss of the cytoplasmic F-actin mesh, associated with spindle migration and cytokinesis defects. In contrast, cofilin inhibition in MII oocytes resulted in a shutdown of actin filament dynamics, together with a considerable expansion of the polarized actin cap. Remarkably, ectopic microvilli outgrowth, restricted to the polarized cortex, was achieved by promoting actin cap disassembly. These data establish cofilin as a critical component in actin network homeostasis in oocytes, in line with cell cycle progression. Furthermore, our findings highlight a role for cofilin in regulating the length of microvillar core bundles, and reveal a mechanism for microvilli elongation based on the disassembly of a competing network.

## Results

### Cofilin activation promotes spindle migration and polar body extrusion

Regulation of cofilin activity is mainly achieved via inhibitory phosphorylation at Ser3, catalyzed by LIMK. While LIMK expression and activation are detected across meiotic maturation in mouse oocytes (Li et al., 2017; Duan et al., 2018), the time course of cofilin phospho-regulation has not yet been described. Thus, we performed western blot analyses using oocyte lysates prepared at four landmark stages of meiotic maturation: prophase arrest (germinal vesicle/GV stage), nuclear envelope breakdown (NEBD), metaphase-I (NEBD+6h; MI) and metaphase-II (MII). Phospho-cofilin was readily detected in GV oocytes, and its level decreased by ∼ 50% at meiosis resumption (NEBD), and decreased further to < 20% of GV level in MI and MII oocytes (Figure 1A), indicating cofilin activation. The drop in phospho-cofilin level was not due to cofilin downregulation, as total cofilin remained unchanged (Figure 1A).

**Figure 1.**
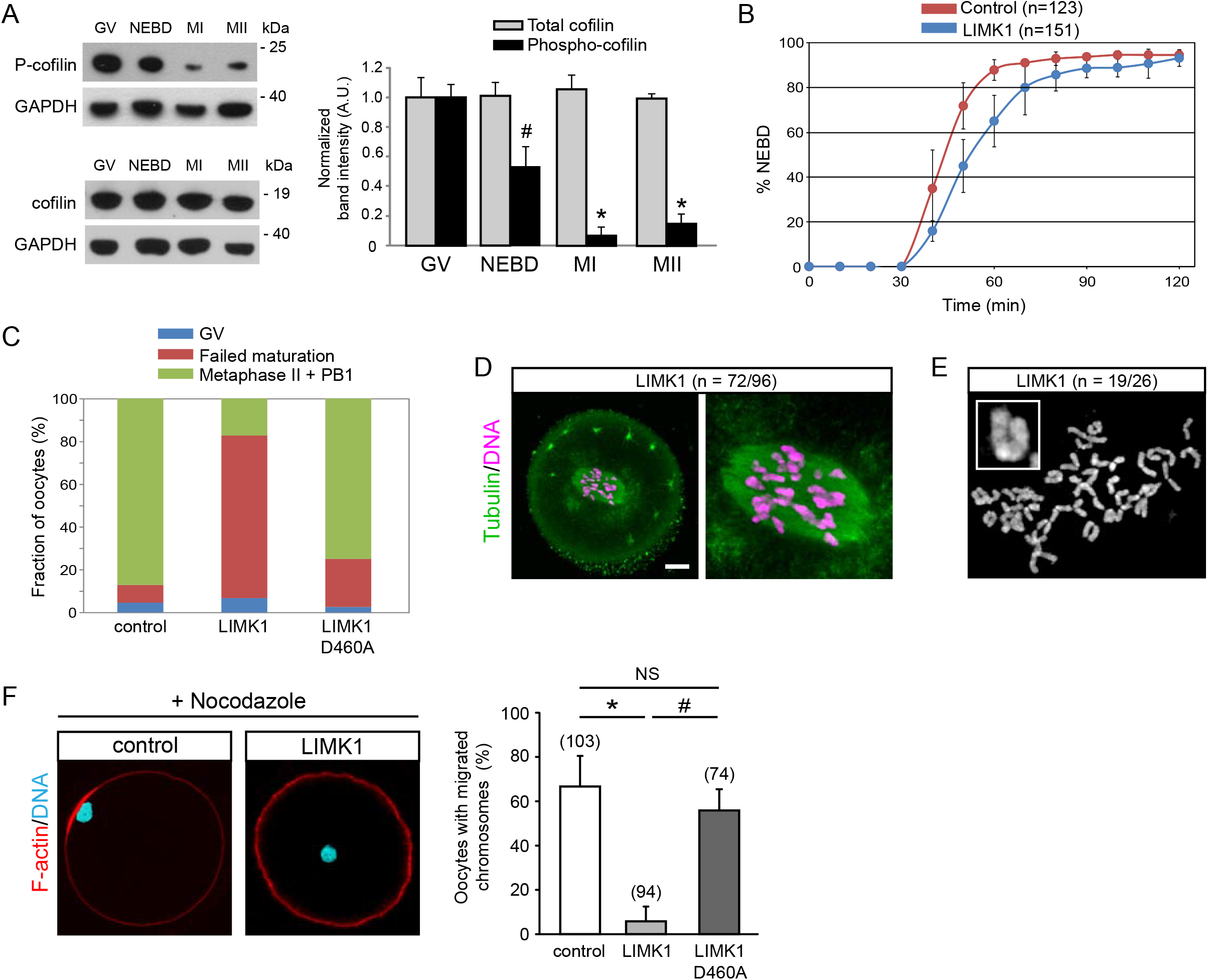
Cofilin activation promotes oocyte maturation. (A) Detection of phospho-cofilin (P-cofilin; top left panel) and total cofilin (bottom left) in oocyte lysates at distinct stages of maturation. GAPDH was used as a loading control. The bar graph (right panel) shows the quantification of band intensities using the GV stage as a reference, and after normalization to the GAPDH signal. Data are means +/- SE of three independent experiments. # : P=0.026 against GV, * : P<10^−4^ against GV. (B) Time course of spontaneous meiosis resumption in oocytes injected with water (Control), or LIMK1 cRNA (LIMK1). Oocytes were injected at the GV stage and cultured for three hours before milrinone washout (t=0). The percentage of oocytes undergoing NEBD was monitored every 10 min for two hours. Data are means +/- SD of four independent experiments. The total number of oocytes scored is indicated in parentheses. (C) Maturation rate of control, LIMK1- and LIMK1^D460A^-expressing oocytes. Oocytes were injected at the GV stage and cultured for three hours before milrinone washout. After overnight culture, oocytes were fixed and stained for spindle microtubules and chromatin, for staging. Oocytes that achieved NEBD but failed to emit the first polar body (PB1) are referred to as “failed maturation”. Over 100 oocytes (3-5 experiments) were scored for each experimental condition. (D) Configuration of the meiotic spindle in a fixed LIMK1-injected oocyte that failed to emit PB1 after overnight culture. Green: anti-tubulin staining; magenta: DNA staining with TO-PRO-3. The right panel shows an expanded view of the spindle. Both images are Z-projections of 13 consecutive confocal frames. Scale bar: 10 µm. (E) Chromosome spread of a LIMK1-injected oocyte that failed to emit PB1 after overnight culture. Note the presence of 40 univalents, indicative of homolog separation. The inset shows an expanded view of a univalent. (F) Rate of chromosome migration to the cortex in nocodazole (5 µM)-treated oocytes. Representative images of a control oocyte and a LIMK1-injected oocyte are shown (left panel). Note the cortical apposition of the chromatin mass, associated with the formation of an actin cap, in control. Red: F-actin stained with Alexa Fluor 568-phalloidin; Cyan: DNA stained with TO-PRO-3. The bar graph (right panel) shows the rate of chromosome migration in nocodazole-treated control, LIMK1- and LIMK1^D460A^-oocytes fixed and stained after overnight culture. Data are means +/- SD of three to five independent experiments. The total number of oocytes scored is indicated in parentheses. *: P<10^−4^, #: P<10^−3^, NS : not significantly different.

To explore the functional significance of cofilin activation, we first attempted cofilin depletion via siRNA injection. However, despite successful knockdown of cofilin mRNA, protein levels were little affected (∼20% depletion; Figure S1), suggesting that cofilin is a relatively abundant and stable protein, a recurrent pitfall in RNA interference strategies in oocytes (Pfender et al., 2015). Alternatively, LIMK1 overexpression is a well established and recognised strategy for acute cofilin inactivation in a variety of cell types (Arber et al., 1998; Yang et al., 1998; Sotiropoulos et al., 1999; Bierne et al., 2001; Takahashi et al., 2001; Amano et al., 2002; Higashida et al., 2013). Hence, we injected prophase oocytes with a cRNA encoding mouse LIMK1, and assessed phospho-cofilin levels by western blot. Surprisingly, phospho-cofilin was not increased further in LIMK1-expressing GV oocytes (Figure S2, left), indicating that cofilin is already largely inactivated during prophase arrest. In contrast, a substantial increase in phospho-cofilin was achieved when LIMK1-injected oocytes were released from prophase arrest and cultured to the MI stage, while the catalytically inactive LIMK1^D460A^ (Yang et al., 1998) had no effect (Figure S2, middle). Spontaneous meiosis resumption was not overtly affected by LIMK1 overexpression (Figure 1B). However, only 20% of LIMK1-injected oocytes had progressed to the MII stage and emitted a polar body after overnight culture, against 87% of control oocytes and 76% of LIMK1^D460A^-injected oocytes (Figure 1C). The majority (72/96 oocytes) of arrested LIMK1-injected oocytes exhibited a bipolar spindle located centrally, pointing to a defect in spindle migration (Figure 1D). Yet, chromosome spreading revealed that most of these arrested oocytes contained 40 univalents (19/26 oocytes examined; Figure 1E), demonstrating they had achieved homologous segregation, but failed to form a polar body.

To support a spindle migration defect, oocytes released from prophase arrest were treated with nocodazole to prevent spindle formation, allowing us to monitor chromosome migration powered by cytoplasmic actin dynamics (Van Blerkom and Bell, 1986; Li et al., 2008). As expected, chromosomes migrated to the cortex in a majority of control and LIMK1^D460A^-injected oocytes, leading to the formation of an actin cap (Figure 1F). In contrast, chromosomes failed to migrate in oocytes expressing LIMK1 (Figure 1F), likely reflecting a defect in actin dynamics.

Altogether, these data show that successful oocyte maturation relies on a switch-like activation of cofilin at meiosis resumption, in order to execute two major actin-dependent processes: spindle migration and polar body formation. Since a centrally-located spindle does not preclude oocyte cleavage at anaphase-I (Pfender et al., 2011; Wei et al., 2018), our data suggest a role for cofilin in cytokinesis, in line with previous findings in cultured somatic cells and Xenopus oocytes (Abe et al., 1996; Hotulainen et al., 2005).

### Cofilin does not contribute to cytoplasmic F-actin dynamics in prophase oocytes

In line with its canonical role in promoting pointed end disassembly, cofilin was suggested to regulate cytoplasmic F-actin dynamics in mouse oocytes (Jang et al., 2014; Montaville et al., 2014; Jo et al., 2016). However, our finding of cofilin being largely inactivated in GV oocytes is inconsistent with this view. To exclude any contribution from residual active cofilin, we overexpressed LIMK1 in prophase-arrested GV oocytes and examined the integrity of the cytoplasmic F-actin mesh using phalloidin staining in fixed oocytes, and EGFP-UtrCH as a probe for F-actin dynamics in live oocytes (Azoury et al., 2008; Schuh and Ellenberg, 2008). Remarkably, LIMK1-injected GV oocytes exhibited a dense cytoplasmic actin network indistinguishable from controls, and showing similarly fast dynamics (Figure 2A, middle and right panels; Movie 1). Moreover, and in line with previous observations (Azoury et al., 2011), LIMK1-injected GV oocytes experienced a significant drop in cytoplasmic F-actin mesh density shortly before NEBD, in the same fashion as control oocytes (Figure 2B; Movie 1). These observations argue against a significant role of cofilin in promoting cytoplasmic F-actin dynamics during prophase arrest in fully grown oocytes. Surprisingly, cofilin also appears dispensable for the collapsing of the actin mesh at meiosis resumption. However, we consistently observed bulky F-actin structures in the nucleoplasm of LIMK1-expressing GV oocytes (see Figure 2A, middle), suggesting that a small pool of active cofilin is engaged in regulating nuclear F-actin turnover during prophase arrest. This is consistent with the reported enrichment of cofilin in the GV nucleoplasm (Jin et al., 2020).

**Figure 2.**
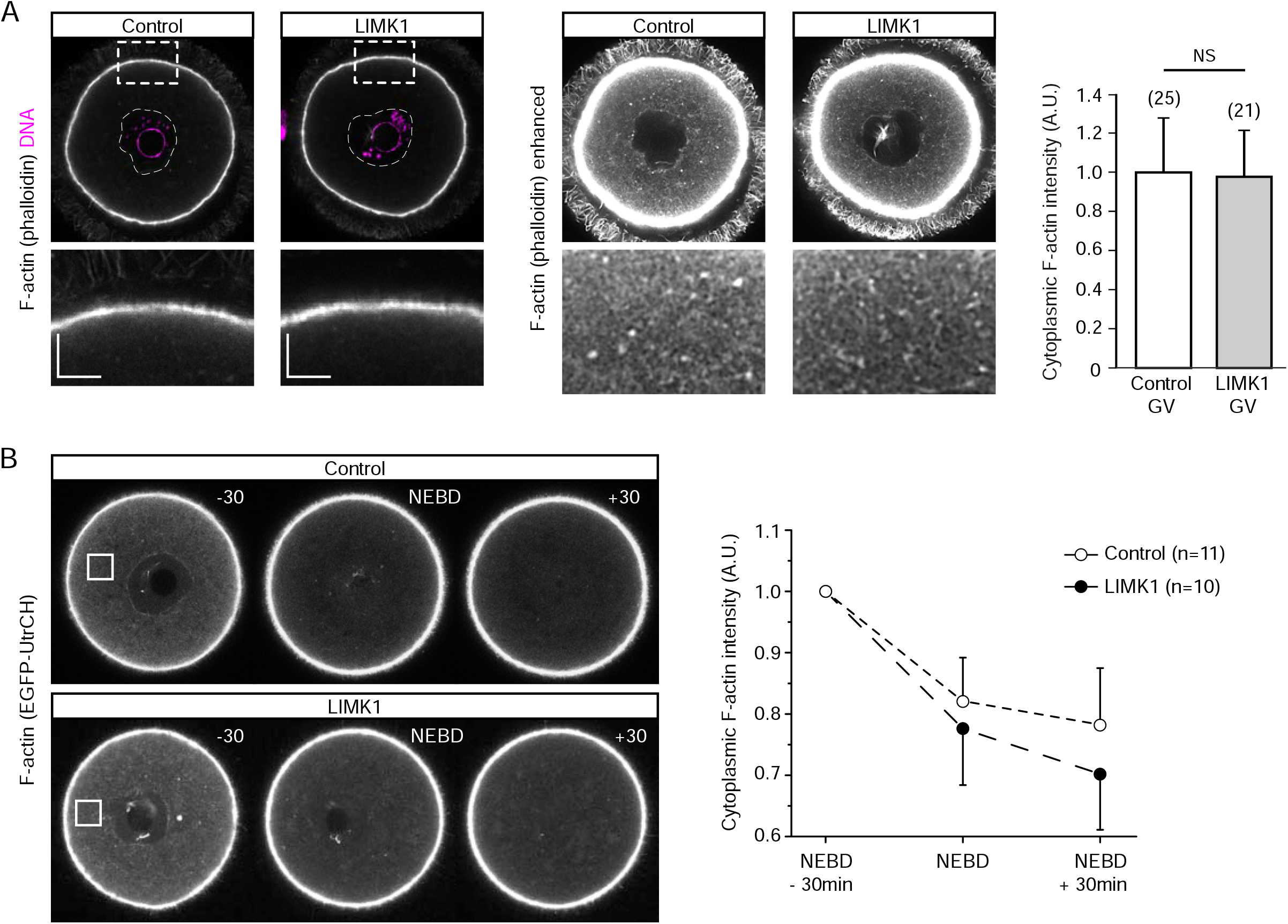
Cofilin is dispensable for F-actin dynamics in prophase oocytes. (A) F-actin staining with Alexa Fluor 568-phalloidin in GV stage oocytes arrested in prophase with milrinone. LIMK1 was overexpressed via cRNA injection followed by culture for 3 hours (LIMK1). Left panels: F-actin imaging with regular settings, avoiding saturation. The GV is depicted by a dashed line. Chromatin is labeled with To-Pro-3 (DNA, magenta). The bottom row shows expanded views of the cortical regions delineated by the dashed boxes. Scale bars: 5 µm. Middle panels: same images with enhanced brightness for better visualization of the cytoplasmic F-actin mesh. The bottom row shows expanded views of a cytoplasmic region (20×15 µm). Note the thick F-actin bundles in the nucleoplasm of the LIMK1-expressing GV oocyte. Right panel: quantification of cytoplasmic F-actin density. The intensity of the phalloidin signal was measured in a cytoplasmic 10×10 µm region. The mean value obtained in control oocytes was normalised to 1. The total number of oocytes examined is indicated in parentheses. Data are means +/- SD of three independent experiments. NS : not significantly different. (B) Live imaging of F-actin networks in control GV (top panel) and LIMK1-injected GV (bottom panel) oocytes expressing EGFP-UtrCH and undergoing meiosis resumption. Still images corresponding to NEBD-30 min, NEBD and NEBD+30min are shown. Fluorescence from the F-actin probe was measured in a 10×10 µm region (white boxes) and plotted against time (means +/- SD; right panel). The values obtained at NEBD-30min were normalized to 1. The number of oocytes examined is indicated in parentheses.

### Cofilin inhibition disrupts cytoplasmic and cortical F-actin networks

In view of the spindle migration defect in LIMK1-injected oocytes, we examined the integrity of the cortical and cytoplasmic F-actin networks, which are both required for successful symmetry breaking (Azoury et al., 2008; Schuh and Ellenberg, 2008; Chaigne et al., 2013). Intriguingly, the cortex of MI oocytes (NEBD+6h) overexpressing LIMK1 was lined with short (1-3 µm in length) linear actin bundles pointing toward both the extracellular milieu and the cell interior (Figure 3A, LIMK1), suggestive of an elongation of microvillar core bundles and cytoplasmic actin rootlets (Wong et al., 1997). Under our experimental conditions, regular microvilli in control oocytes could not be resolved accurately (Figure 3A, Control), due to their fairly short length (<0.5 µm; Benammar et al., 2017). Interestingly, while the cytoplasmic F-actin network was readily detectable in control MI oocytes, it had virtually vanished in LIMK1-expressing MI oocytes (Figure 3B,C, LIMK1). Live imaging of F-actin dynamics confirmed the absence of the cytoplasmic actin mesh in LIMK1-expressing MI oocytes (Movie 2). In contrast, expression of catalytically-inactive LIMK1^D460A^ did not alter microvilli length nor the integrity of the cytoplasmic F-actin mesh (Movie 2), indicating that the observed F-actin defects were effectively linked to the kinase activity of LIMK1. To confirm the specificity of the phenotype, oocytes were co-injected with LIMK1 and constitutively active cofilin XAC(A3) (Takahashi et al., 2001) in an attempt to rescue F-actin severing activity. Expression of the non-phosphorylatable mutant cofilin effectively abolished microvilli overgrowth in LIMK1-expressing oocytes (Figure 3A,B and Figure 4C,D). Cytoplasmic F-actin staining did not recover however (Figure 3B,C), consistent with the previous observation that constitutively active cofilin dissolves the F-actin mesh (Jin et al., 2020).

**Figure 3.**
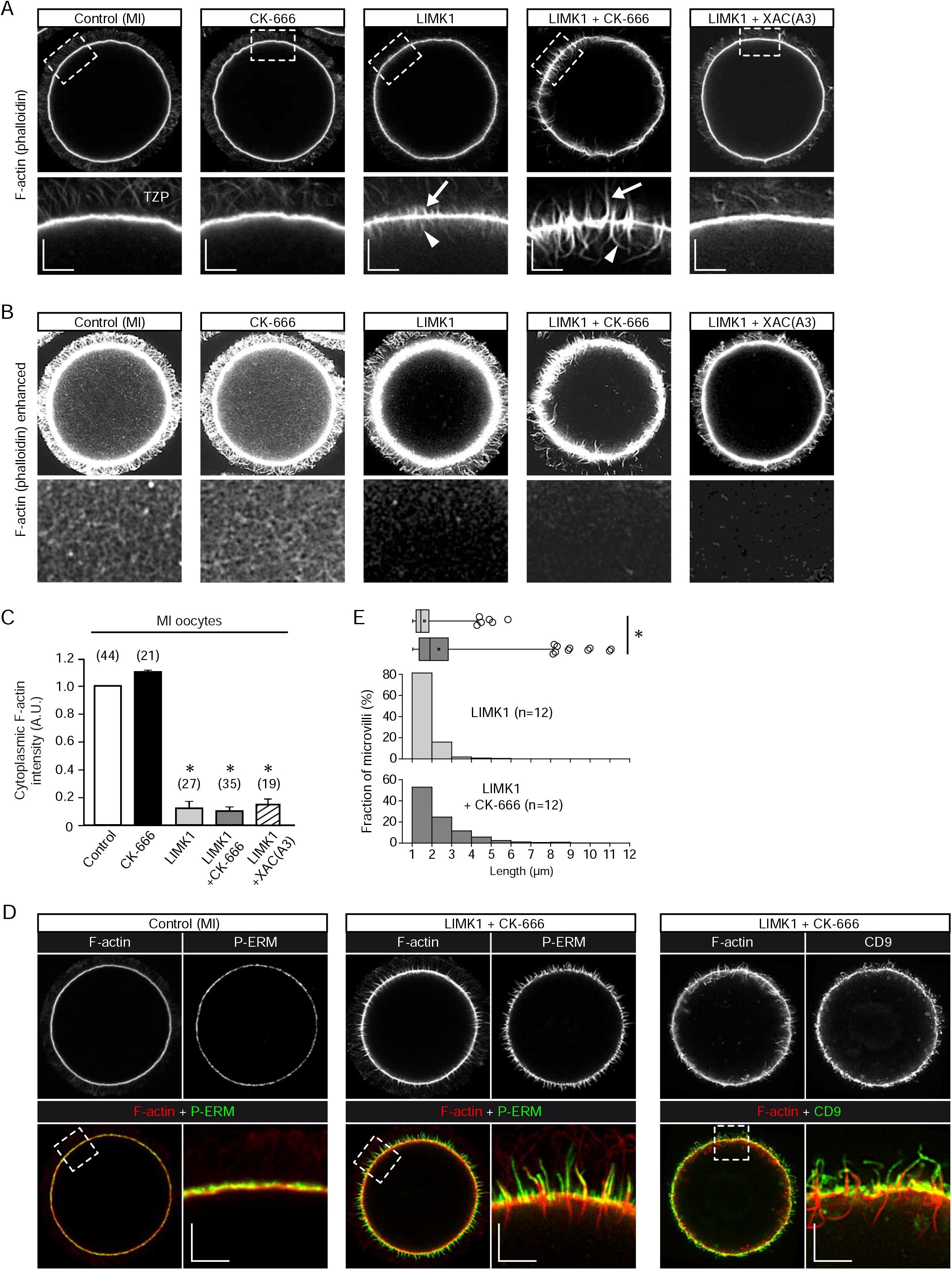
Cofilin regulates cortical and cytoplasmic F-actin homeostasis in MI oocytes. (A) F-actin staining with Alexa Fluor 568-phalloidin in fixed MI (NEBD+6h) oocytes, under various experimental conditions. LIMK1 cRNA was injected at the GV stage followed by in vitro culture to NEBD+6h (LIMK1). CK-666 (100 µM) was added at the time of milrinone washout to study the effects of Arp2/3 inhibition alone (CK-666), or in combination with LIMK1 overexpression (LIMK1 + CK-666). Coexpression of LIMK1 and constitutively-active cofilin XAC(A3) was achieved by coinjecting the respective cRNAs in GV oocytes (LIMK1 + XAC(A3)). The bottom panels are expanded views of the cortical regions delineated by the dashed boxes. The arrows and arrowheads point to elongated microvillar actin bundles, and cytoplasmic actin rootlets, respectively. Note that transzonal projections, which originate from surrounding follicular cells, are also faintly labeled (TZP). The scale bars represent 5 µm. (B) Same images as top row in (A), with enhanced brightness for better visualization of the cytoplasmic F-actin mesh (or absence of). The bottom panels are expanded views of a cytoplasmic region (20×15 µm) avoiding the spindle area and actin rootlets. (C) Quantification of cytoplasmic F-actin in fixed MI oocytes, as shown in (B). The mean intensity of the phalloidin signal was measured in a cytoplasmic 10×10 µm region. The mean value obtained in control oocytes was normalised to 1. The number of oocytes examined is indicated in parentheses. Data are means +/- SD of three to five independent experiments. *: P<10^−4^ against Control. (D) Control MI (left panel) and LIMK1-expressing MI oocytes treated with 100 µM CK-666 (middle and right panels) were fixed and stained for F-actin (Alexa Fluor 568-phalloidin; red in the merge image) together with P-ERM (left and middle; green) or CD9 (right; green). Expanded views of the cortical regions (dashed boxes in the merge images) are shown. See also Movie 4. (E) Effect of Arp2/3 inhibition on microvilli length. LIMK1-expressing oocytes were fixed at NEBD+6h and microvilli were detected using FiloQuant (with a cut-off at 1 µm) in a single confocal section taken at the equator of the cell. Microvilli lengths were pooled into 1-µm size bins plotted against frequency. Corresponding box plots (same *x* axis) are shown at the top, with higher values displayed as outliers. Twelve oocytes were analyzed in each condition. *: P<10^−4^ from Mann-Whitney test.

**Figure 4.**
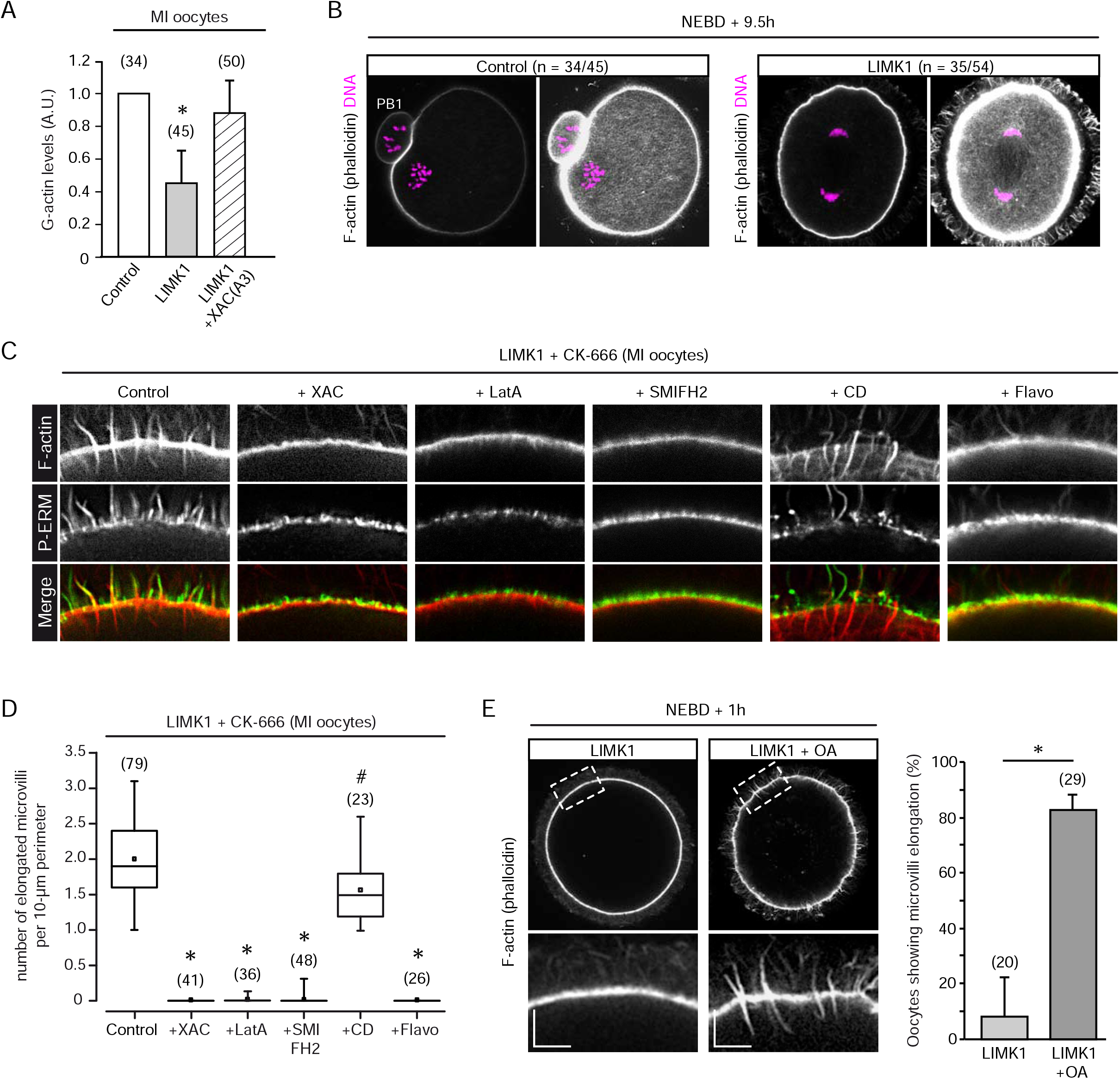
Microvilli elongation requires G-actin, formin and CDK1 activities. (A) G-actin levels in MI oocytes. Control oocytes, or oocytes expressing LIMK1 alone or in combination with XAC(A3) were fixed at MI (NEBD + 6h) and processed for DNAse 1 staining to label G-actin. The number of oocytes examined is indicated in parentheses. Data are means +/- SD from four independent experiments. *: P<10^−4^ against Control. (B) Chromosome (DNA; To-Pro-3, magenta) and F-actin (Alexa Fluor 568-phalloidin) staining in a control oocyte shortly after polar body extrusion (left panel), and an oocyte injected with LIMK1 cRNA at the MI (NEBD + 6h) stage (right panel). In both cases, oocytes were fixed at NEBD + 9.5h. Note the anaphase-I configuration of the LIMK1-expressing oocyte. In both panels, the left image shows F-actin staining with regular settings, and the right image shows the same F-actin staining with enhanced brightness for better visualization of the cytoplasmic F-actin mesh. The number of oocytes examined showing a similar pattern is indicated in parentheses. PB1 : first polar body. (C) Effects of various treatments on microvilli elongation. All images are expanded views of the cortical region of LIMK1-expressing MI oocytes treated with CK-666, stained for F-actin (Alexa Fluor 568-phalloidin, red) and P-ERM (green). (D) Quantification of the density of elongated microvilli under conditions shown in (C). Elongated microvilli were counted along the oocyte perimeter, in a single confocal image taken at the equator of the oocyte, and expressed as a density per 10-µm perimeter. The number of oocytes examined for each experimental condition is indicated in parentheses. XAC: constitutively active XAC (A3); LatA: latrunculin A (0.3 µM); SMIFH2: SMIFH2 (30 µM); CD: cytochalasin D (100 nM); Flavo: flavopiridol (5 µM). Statistical significance (Student’s *t*-test) was calculated against the control condition (LIMK1+CK-666). *: P<10^−4^, # : P=0.001. (E) F-actin staining (Alexa Fluor 568-phalloidin) in LIMK1-expressing oocytes treated with DMSO (LIMK1, left) or 2.5 µM okadaic acid (LIMK1 + OA, right). Oocytes were injected with LIMK1 cRNA at the GV stage, and cultured in vitro to NEBD+1h before fixation. Lower panels are expanded views of the cortical regions delineated by the dashed boxes. The bar graph shows the percentage of oocytes showing microvilli elongation. Data are means +/- SD from three independent experiments. The number of oocytes examined is indicated in parentheses. *: P=0.001. Scale bars in (E) represent 5 µm.

To test whether microvillar actin overgrowth was related to the reported Arp2/3-dependent cortical actin thickening (Chaigne et al., 2013), we supplemented the culture medium with the Arp2/3 inhibitor CK-666 (Nolen et al., 2009). Strikingly, Arp2/3 inhibition exacerbated microvillar actin outgrowth, in the form of thick bundles intersecting the cortical actin layer, tapered at both ends and often heavily bent (Figure 3A, LIMK1+CK-666; Movie 3). Accordingly, outward processes were decorated by phosphorylated Ezrin/Radixin/Moesin (P-ERM; Figure 3D and Movie 4) and the tetraspanin CD9 (Figure 3D), two classic markers of microvillar identity (Runge et al., 2007; Pelaseyed and Bretscher, 2018). To quantify the effects of CK-666 on microvilli length, we used Filoquant, an ImageJ plugin originally designed to extract filopodia parameters (Jacquemet et al., 2017; Figure S3). While detected microvilli were mostly in the 1-2 µm range in LIMK1-overexpressing MI oocytes, CK-666 shifted their length toward the 2-5 µm range, with some microvilli reaching over 10 µm (Figure 3E). These values are likely underestimated as a number of “broken” microvilli could not be resolved along their entire length within the plane of the confocal section (see Figure S3). Importantly, microvilli overgrowth, and the concurrent loss of cytoplasmic F-actin, were not observed in oocytes treated with CK-666 alone (Figure 3A-C, CK-666). In contrast, microvilli length occasionally reached 4-6 µm in LIMK1-injected MI oocytes without CK-666 addition (Figure 3E), indicating that Arp2/3 inhibition was synergistic, but not an absolute requirement.

Collectively, these results show a requirement for cofilin activation at meiosis resumption, in order to preserve the integrity of the cortical and cytoplasmic F-actin networks of mouse oocytes. Interfering with cofilin activation leads to microvilli overgrowth and a reciprocal loss of cytoplasmic F-actin, suggesting that microvilli acted as a sink for actin monomers, at the expense of F-actin mesh reassembly during maturation.

### Microvillar actin elongation relies on G-actin supply, CDK1 and formins

In view of the unaltered F-actin mesh disassembly at meiosis resumption in cofilin-inhibited oocytes (see Figure 2B), we speculated that the released actin monomers were diverted to microvilli, precluding cytoplasmic mesh reassembly later during maturation. Consistent with this idea, LIMK1-expressing MI oocytes showed significantly lower levels of cytoplasmic G-actin, as visualized by DNAse I staining (Figure 4A). Accordingly, co-expression of XAC(A3) to prevent microvilli overgrowth restored the G-actin pool to near-control levels (Figure 4A), consistent with microvilli acting as a sink for monomers. To investigate further a possible link between the collapsing of the F-actin mesh and microvillar elongation, we injected LIMK1 cRNA after oocytes had reached MI (NEBD+6h) and recovered a dense cytoplasmic F-actin mesh (as shown in Figure 3B, Control). After additional culture for 3 hours to allow for protein expression (NEBD+9.5h), oocytes were fixed and stained for chromosomes and F-actin. As illustrated in Figure 4B, a large fraction (35/54 oocytes; 65%) of these oocytes was blocked in an anaphase-I configuration, while the majority of time-matched control oocytes had emitted the first polar body (34/45 oocytes, 76%). This is consistent with the cytokinesis failure described earlier for oocytes injected at the GV stage (Figure 1C-E). Interestingly, these anaphase I-arrested oocytes showed no sign of microvilli elongation (0/54 oocytes), and exhibited a dense cytoplasmic F-actin network akin to control oocytes (Figure 4B). These findings indicate that in oocytes with an intact cytoplasmic mesh, LIMK1 overexpression is not sufficient to induce microvillar actin overgrowth. This is consistent with the observation that GV oocytes, in which cofilin is largely inhibited, do not show microvillar overgrowth, and retain a dense actin mesh.

Consistent with a requirement for actin monomer supply, sequestration of G-actin with Latrunculin A (0.3 µM; Coué et al., 1987) prevented microvilli overgrowth in LIMK1-expressing MI oocytes, suggesting that microvillar actin elongation proceeds via monomer addition at filament ends (Figure 4C,D; +LatA). We next tested the effect of a low dose of cytochalasin D (CD; 100 nM), in order to achieve barbed end capping (Sampath and Pollard, 1991). However, microvilli elongation was largely resistant to CD treatment (Figure 4C,D; +CD), arguing that barbed ends may be protected against capping. In this regard, formins exhibit anti-capping activity due to their persistent association with the growing barbed end (Chesarone et al., 2010; Shekhar et al., 2016). Accordingly, the pan-formin inhibitor SMIFH2 (30 µM; Rizvi et al., 2009) abolished microvilli outgrowth, while regular microvilli were preserved (Figure 4C,D; +SMIFH2). In our search for a candidate formin, we observed that DAAM1, which is constitutively enriched at the oocyte cortex (Lu et al., 2017), accumulated at the distal tips of enlarged microvilli (Figure S4A), possibly reflecting a role in filament elongation. DAAM1 also decorated the entire length of microvillar core bundles and actin rootlets (Figure S4A), in line with its reported role in filament bundling (Jaiswal et al., 2013). In contrast, Fmn2, which is also enriched at the oocyte cortex (Pfender et al., 2011), did not accumulate along the length nor the tips of elongated microvilli (Figure S4B).

The fact that prophase oocytes do not show excessive microvilli nor actin rootlet elongation (Figure 2A), despite a substantial phosphorylation of their cofilin pool (Figure S2), is suggestive of an additional regulatory mechanism related to cell cycle resumption. To test this idea, LIMK1-injected GV oocytes arrested in prophase by milrinone were treated with the PP1/PP2A inhibitor okadaic acid (OA, 2.5 µM) in order to accelerate substrate phosphorylation by CDK1, thereby forcing oocytes into meiosis resumption (Rime et Ozon, 1990). Remarkably, OA treatment accelerated microvilli and actin rootlet overgrowth, which became detectable as early as one hour after NEBD, at which time vehicle-treated LIMK1-expressing oocytes did not show yet any sign of overgrowth (Figure 4E). To establish a specific requirement for CDK1, the CDK1 inhibitor flavopiridol (5 µM; Wei et al., 2018) was added to the culture medium one hour after NEBD and oocytes were further cultured until MI (NEBD+6h). CDK1 inhibition effectively abrogated microvillar actin overgrowth in LIMK1-expressing oocytes (Figure 4C,D; +Flavo). Regular microvilli were still detectable however, suggesting that the lack of microvilli elongation was not due to an intrinsic defect in microvillar integrity.

Taken together, these data suggest that microvilli overgrowth relies on a supply of free actin monomers, to which the collapse of the F-actin mesh is likely to contribute. However, the microvillar elongation phenotype is not the mere consequence of cofilin inactivation. It also relies on CDK1 substrate phosphorylation, and may involve a formin-dependent elongation machinery. This outgrowth mechanism appears to build upon regular microvilli, which seem otherwise unaffected by CDK1 or formin inhibition.

### Cofilin controls F-actin dynamics and the size of the actin cap in MII oocytes

We next explored the role of cofilin in regulating steady-state F-actin dynamics in oocytes that have achieved maturation and arrested at MII. Freshly ovulated MII oocytes were injected with LIMK1 cRNA for acute cofilin inhibition, as reflected in an increase in the phospho-cofilin signal (Figure S2, right). After 2-3 hours of culture, LIMK1-expressing MII oocytes exhibited a dramatic overgrowth of the actin cap, reaching up to 15 µm-deep into the inner cytoplasm (Figure 5A-C). Remarkably, cytoplasmic material was excluded from this actin-filled volume, consistent with a dense network with a small mesh size (Figure 5B, asterisk). While cortical actin was moderately thickened in the rest of the cortex, there was no sign of microvilli overgrowth (Figure 5B,D). In contrast, catalytically-inactive LIMK1^D460A^ did not alter cortical actin dynamics, in line with the lack of cofilin phosphorylation (Figure S2 and 5C,D).

**Figure 5.**
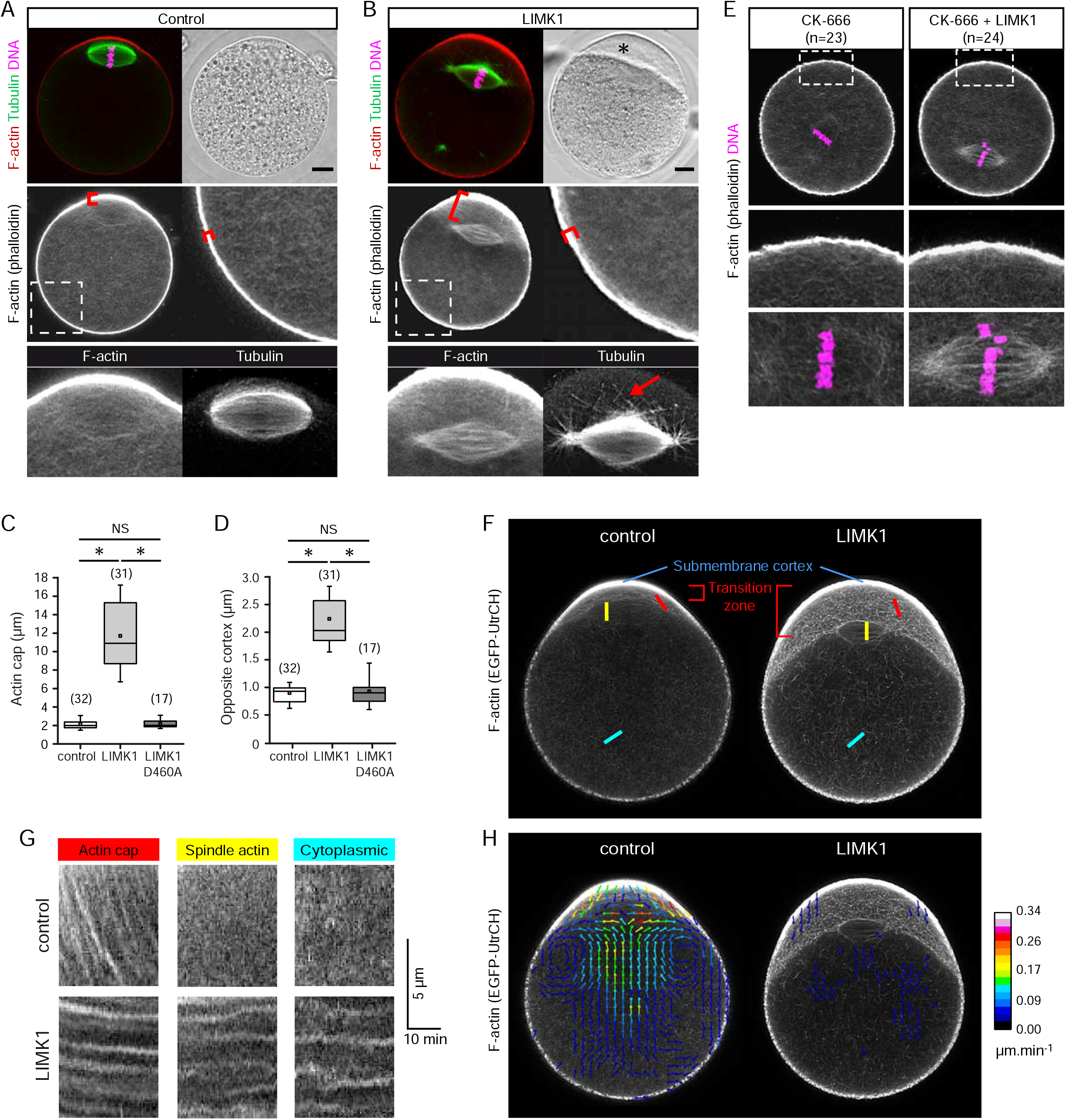
Cofilin controls actin network dynamics in MII oocytes. (A,B) Staining of F-actin and spindle microtubules in a control MII oocyte (A) and a LIMK1-injected MII oocyte (B). Upper panels show the brightfield and merge fluorescence images. Note the exclusion of cytoplasmic material in the region of actin cap expansion (B; asterisk). Red: F-actin (Alexa Fluor 568-phalloidin); green: anti-tubulin; magenta: chromosomes stained with TO-PRO-3 (DNA). Scale bars represent 10 µm. Middle panels show the F-actin channel with enhanced brightness for better visualization of cytoplasmic and spindle F-actin. Magnified views of the non polarized cortex (dashed boxes) are shown. Lower panels are expanded views of the spindle region. Note the increased spindle actin signal and the elongated microtubules (arrow) projecting toward the expanded actin cap in the LIMK1-expressing oocyte (B). (C,D) Quantification of the thickness of the actin cap (C) and the opposite cortex (D), as illustrated by the red brackets in (A,B). The number of oocytes examined is indicated in parentheses. *: P<10^−5^, NS: not significantly different. (E) Staining of F-actin networks (Alexa Fluor 568-phalloidin) in MII oocytes treated with CK-666 (100 µM, 4h) alone (left), or CK-666 followed by LIMK1 cRNA injection (right). Middle and bottom panels are expanded views of the cortical area (dashed boxes) and spindle area, respectively. Note the increased spindle actin signal, but the lack of microvilli elongation, in the oocyte expressing LIMK1. The total number of oocytes examined is indicated in parentheses. (F) Visualization of F-actin networks in live MII oocytes expressing EGFP-UtrCH alone (left) or together with LIMK1 (right). The polarized actin cap is defined as a submembrane cortical layer of high intensity (submembrane cortex) and a transition zone of milder intensity corresponding to actin filaments flowing away from the cap. See also Movie 5. (G) Kymographic analysis of actin flow in the actin cap transition zone, spindle and cytoplasm, as depicted by the colored regions in (F). (H) STICS analysis of F-actin dynamics in the oocytes shown in (F). The color-coded vector maps were generated from the time-lapse recordings shown in Movie 5. The heat bar indicates filament velocity (µm.min^-1^). Data in (F-H) are representative of 10 (control) and 8 (LIMK1) similar observations.

As a consequence of actin cap expansion, the MII spindle was distanced from the cortex, yet we never observed spindle detachment (Figure 5B). Intriguingly, elongated spindle microtubules appeared to infiltrate the oversized actin cap, possibly contributing to a spindle-actin anchoring mechanism (Figure 5B, arrow). The spindles also appeared highly focused at the poles (Figure 5B), a feature previously reported in spindles artificially enriched in actin (Mogessie and Schuh, 2017). Accordingly, while spindle actin generated a faint signal in control oocytes, it was readily observable in LIMK1-injected MII oocytes, likely reflecting filament stabilization and/or thickening (Figure 5A,B, lower panels).

We next tested whether the absence of microvilli overgrowth was due to actin monomers being diverted to the expanding actin cap. To this end, MII oocytes were injected with LIMK1 cRNA after prior treatment with CK-666 to inhibit Arp2/3. Consistent with previous observations (Yi et al., 2011), prolonged incubation with CK-666 (100 µM, 4h) induced a loss of the actin cap, along with spindle detachment from the cortex (Figure 5E, left panels). Subsequent LIMK1 expression still failed to induce microvilli elongation (0/24 oocytes), while spindle actin was markedly increased, and the cytoplasmic mesh was preserved (Figure 5E, right panels). Therefore, the absence of microvilli elongation cannot be attributed to monomer sequestration into the oversized actin cap.

In order to appreciate the effects of cofilin inactivation on actin network dynamics, we monitored actin fluxes in live MII oocytes expressing EGFP-UtrCH. Consistent with previous studies (Yi et al., 2011; Mogessie and Schuh, 2017), control oocytes exhibited highly dynamic cytoplasmic and spindle actin networks, as well as a retrograde actin flow originating from the polarized cap (Movie 5). This retrograde actin flow, visualized as a transition zone of milder intensity pervading the subcortical cytoplasm, was readily captured by kymographic analysis, while cytoplasmic and spindle actin dynamics were too fast to be resolved (Figure 5F,G). However, all three actin networks were successfully captured and quantified by spatiotemporal image correlation spectroscopy (STICS) analysis (Yi et al., 2011), revealing highest velocities in the actin cap transition zone and spindle actin (Figure 5H). In stark contrast, actin networks were virtually static in LIMK1-injected MII oocytes (Movie 5), as reflected in the flat lines on the kymographs, and a faint STICS signal (Figure 5G,H). Live imaging also revealed that actin cap overgrowth resulted from the expansion of the transition zone, which remained segregated from the subjacent actin mesh by a sharp interface (Figure 5F and Movie 5).

Collectively, these data provide three significant informations: 1) cofilin promotes global actin network turnover in mature oocytes, and in particular, restrains actin cap expansion; this is in contrast with the prevailing model by which cofilin amplifies the branching activity of Arp2/3 (Ichetovkin et al., 2002; Kiuchi et al., 2007); 2) in the absence of functional cofilin, the cytoplasmic F-actin network is frozen, in stark contrast with prophase oocytes where it remains highly dynamic, thus reinforcing the idea of distinct regulatory mechanisms in line with the cell cycle stage; and 3) in line with our observations in prophase oocytes and oocytes injected with LIMK1 after reaching the MI stage, cofilin inhibition, per se, is not sufficient to induce microvilli overgrowth in MII oocytes.

### Actin cap disassembly promotes microvilli elongation

We reasoned that MII oocytes may experience microvilli elongation if the G-actin pool was raised to a critical threshold. To test this prediction, we treated MII oocytes with CK-666 (100 µM) for 3 hours such as to promote spontaneous actin cap disassembly and monomer release. Remarkably, CK-666 triggered the outgrowth of a tuft of elongated microvilli in the polarized cortex overlying the actin cap, a region otherwise devoided of microvilli (Figure 6A,B,I). As shown above for LIMK1-expressing MI oocytes, these ectopic microvilli were positive for P-ERM (Figure 6B) and CD9 (Movie 6), and were decorated along their length by DAAM1, which also accumulated at the tips (Figure S4C). Accordingly, SMIFH2 prevented the outgrowth of these ectopic microvilli, while leaving regular microvilli unaffected (Figure 6C,I). Likewise, monomer sequestration with latrunculin A prevented ectopic microvilli elongation (Figure 6D,I). Moreover, while CD (100 nM) effectively collapsed regular microvilli, as evidenced by a substantial loss of P-ERM staining (Figure 6E, arrow), it did not prevent the outgrowth of ectopic microvilli in response to CK-666 (Figure 6E,I). In fact, exposure to CD alone resulted in a similar burst of ectopic microvilli, while the regular microvilli were depleted (Figure 6F,I). As stated above, CD (100 nM) is expected to act as a barbed end capper, suggesting that the collapse of regular microvilli reflected unbalanced pointed end disassembly. Accordingly, injection of LIMK1 cRNA shortly (1h) before CK-666 addition - such as to retain residual cofilin activity for actin cap disassembly - resulted in further elongation of ectopic actin rootlets (Figure 6G). Of note, ectopic microvilli were no longer detected in oocytes where the actin cap had fully disassembled, and the spindle relocated to the oocyte center, by extending the CK-666 treatment to 5 hours (Figure 6H,I). This observation suggests that ectopic microvilli are transient structures relying on actin monomers supplied by the disassembling actin cap. Together, these results suggest that an increase in G-actin supply can elicit microvilli elongation in MII oocytes, with similar regulatory properties as the elongated microvilli depicted in MI oocytes overexpressing LIMK1. Further elongation of ectopic actin rootlets upon LIMK1 expression provides compelling evidence that cofilin regulates the length of microvillar actin filaments in oocytes.

**Figure 6.**
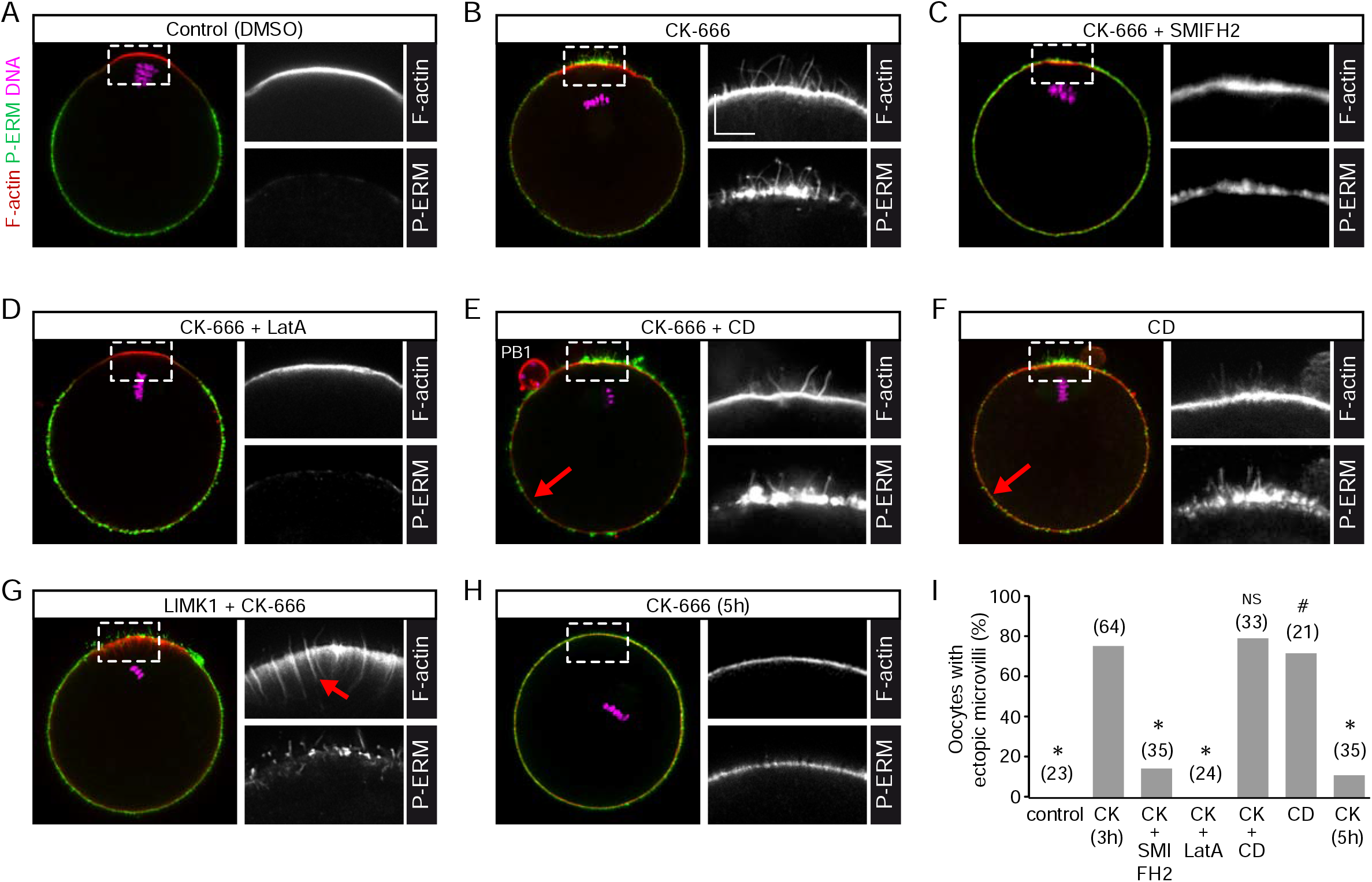
Actin cap disassembly promotes microvilli outgrowth. MII oocytes were fixed and stained for F-actin (Alexa Fluor 568-phalloidin, red), P-ERM (green) and chromosomes (DNA; TO-PRO-3, magenta). Magnified views of the F-actin and P-ERM signals in the polarized cortex (dashed boxes) are shown. (A) Control MII oocyte, treated with vehicle (0.1 % DMSO) for 3 hours. Note the complete absence of microvilli in the polarized cortex. (B) MII oocyte treated with CK-666 (100 µM) for 3 hours. Note the tuft of elongated microvilli in the polarized cortex and the detached MII spindle. (C) MII oocyte treated with CK-666 and SMIFH2 (30 µM) for 3 hours. Note the absence of ectopic microvilli in the polarized cortex and the persistence of regular microvilli in the rest of the cortex. (D) MII oocyte treated with CK-666 (100 µM) and Latrunculin A (LatA; 0.3 µM) for 3 hours. (E) MII oocyte treated with CK-666 (100 µM) and cytochalasin D (CD; 100 nM). Here, treatment was for 2 hours only, as CD accelerated the loss of the actin cap. Note the large gaps in P-ERM staining in the non polarized cortex (arrow), reflecting the collapse of regular microvilli. PB1: first polar body. (F) MII oocyte treated with CD (100 nM) alone, for 1 hour. Note the tuft of microvilli in the polarized cortex, while regular microvilli are depleted (arrow). (G) LIMK1-expressing MII oocyte treated with CK-666 (100 µM) for 3 hours. CK-666 was added 1 hour after LIMK1 cRNA injection in order to retain residual cofilin activity. Note the cytoplasmic actin rootlets extending from the ectopic microvilli (arrow). (H) MII oocyte treated with CK-666 (100 µM) for 5 hours. All images (A-H) are Z-compressions of three consecutive 1 µm-thick confocal sections. The scale bar in (B, F-actin) is 5 µm and applies to all magnified images in this Figure. (I) Rate of ectopic microvilli outgrowth in MII oocytes under the experimental conditions depicted in (A-F,H). The bars indicate the fraction (%) of oocytes showing ectopic microvilli. The number of oocytes examined is indicated in parentheses. * : P<10^−4^ against CK(3h); NS : not significantly different against CK(3h); # : P=0.01 against control.

## Discussion

Adding to the growing repertoire of actin filament regulators in oocytes (Uraji et al., 2018; Duan and Sun, 2019), we show that meiosis resumption involves a switch-like activation of cofilin, that is instrumental in regulating actin network size and dynamics, for successful maturation (Figure 7A). Considering its universal role in actin homeostasis, our finding that cofilin is largely inactivated in GV oocytes is intriguing, and suggests that alternative mechanisms exist to achieve filament turnover during prophase arrest. Likewise, Xenopus oocytes were reported to contain mostly phosphorylated XAC (Abe et al., 1996; Iwase et al., 2013). Cofilin activation at meiosis resumption may involve the upregulation of cofilin phosphatases, such as members of the Slingshot family (Kaji et al., 2003; Iwase et al., 2013).

**Figure 7.**
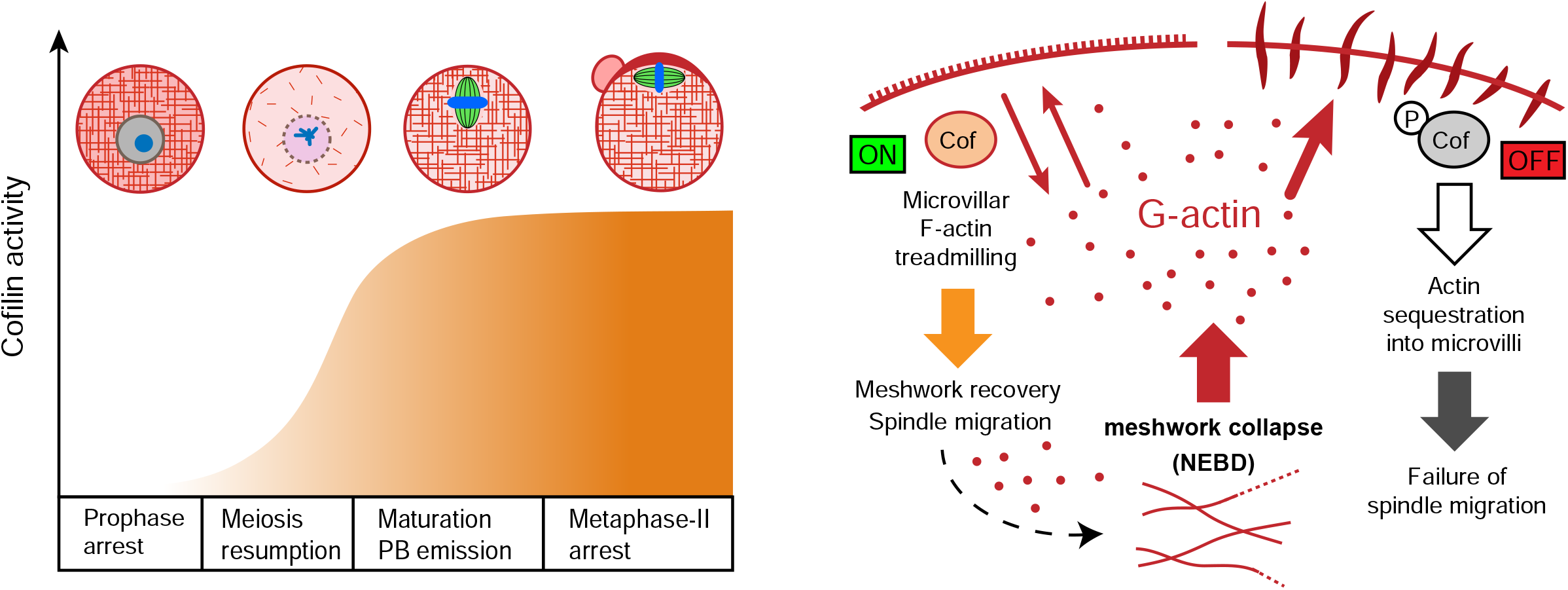
Model depicting the regulation of actin homeostasis by cofilin. Left panel : Theoretical time course of cofilin activation during oocyte maturation. Corresponding meiotic stages are illustrated, with actin filaments in red. PB: polar body. Right panel : model depicting how cofilin inactivation (phosphorylation) leads to microvilli elongation and a loss of the cytoplasmic F-actin mesh. The collapse of the F-actin mesh at meiosis resumption is assumed to release free actin monomers (G-actin, red dots) that are captured by the barbed ends of microvillar actin filaments. Concurrently, cofilin activation (left side; cofilin “ON”) accelerates pointed end disassembly, thereby maintaining the stationary length of microvilli through treadmilling. In doing so, cofilin also maintains the cytoplasmic G-actin pool for recovery of the F-actin mesh later during maturation, enabling spindle migration. Inactivation of cofilin by phosphorylation (right side; cofilin “OFF”) prevents pointed end depolymerization, resulting in exaggerated growth of microvillar core bundles, acting as a sink for monomers. As a result, the actin mesh is not rebuilt and spindle migration fails. Microvilli elongation requires CDK1 and formin activities (not depicted).

In contrast to other actin-rich processes such as lamellipodia and filopodia, which regulatory mechanisms have been extensively described, microvillar actin nucleation and dynamics are still poorly understood (Crawley et al., 2014; Pelaseyed and Bretscher, 2018). Except for their localized disassembly during cortical polarization (Maro et al., 1986; van Blerkom and Bell, 1986; Dehapiot and Halet, 2013), mouse oocyte microvilli exhibit little change in size throughout meiotic maturation (Benammar et al., 2017). Yet, our data demonstrate that they remain dynamic structures amenable to extensive growth in response to actin homeostasis perturbation.

We document a striking elongation of microvilli, and a reciprocal loss of the F-actin mesh, in MI oocytes injected with LIMK1 cRNA at the GV stage to prevent cofilin activation. However, our data in GV- and MII-arrested oocytes demonstrate that cofilin inactivation alone is not sufficient to trigger microvilli elongation, nor F-actin mesh breakdown. This is corroborated by the absence of microvillar elongation or mesh disassembly, when LIMK1 is injected at the MI stage. The microvillar elongation phenotype seems therefore to arise specifically when cofilin is inactivated from meiosis resumption onward. This is precisely the time when the cytoplasmic F-actin mesh experiences a substantial decrease in density, followed by a slow recovery (Azoury et al., 2011; Holubcová et al., 2013; Wei et al., 2018). Thus, one possible scenario is that the release of actin monomers upon mesh disassembly worked as a cue in driving microvillar actin elongation in cofilin-inactivated oocytes. In line with this idea, microvilli elongation consecutive to an increase in G-actin supply was initially reported in isolated brush borders (Mooseker et al., 1982; Stidwill and Burgess, 1986). Furthermore, an increase in G-actin was shown to promote actin filament growth via a direct stimulation of formin activity (Higashida et al., 2013). The synergistic effect of CK-666 may reflect an additional supply of monomers, via the disassembly of an Arp2/3-dependent cortical actin layer (Chaigne et al., 2013). Alternatively, the Arp2/3 complex may restrain microvilli elongation via a direct interaction with a formin machinery, as previously shown for mDia2-dependent filopodia formation (Beli et al., 2008). However, it is unknown whether such inhibitory complexes are disassembled by CK-666. While the formin Daam1 appears as a likely candidate in our system, its role in microvilli elongation remains to be formally established. Besides, off-target effects were recently reported for SMIFH2, hence data using this compound should be interpreted with caution (Isogai et al., 2015; Nishimura et al., 2021. Other types of elongation factors may contribute to the observed microvilli overgrowth, such as IRTKS and EPS8, which role in microvilli elongation was recently characterized in epithelial cells (Postema et al., 2018).

Microvilli outgrowth consecutive to actin cap disassembly in MII oocytes provides further evidence for microvilli length and/or biogenesis being regulated by G-actin availability. Likewise, a recent study documented epithelial microvilli elongation consecutive to Arp2/3 inhibition in differentiating enterocytes (Faust et al., 2019). It is remarkable that ectopic microvilli in mouse oocytes were restricted to the otherwise amicrovillar polarized cortex. Treatment with CK-666 is expected to release actin monomers locally (Burke et al., 2014; Suarez and Kovar, 2016). Hence we suggest that, by preventing monomer recycling into new or existing branches (Vitriol et al., 2015), CK-666 and CD have both contributed to increasing the G-actin pool locally, leading to microvilli outgrowth in the nearby cortex. In turn, it is tempting to assume that the regulated disassembly of microvilli during the process of oocyte polarization, proceeds via a competition for monomers between the Arp2/3 complex and the microvillar machinery, at the expense of the latter. The growth of corresponding ectopic actin rootlets after additional LIMK1 expression provides clear evidence for cofilin regulating microvilli length in oocytes.

We therefore propose that the switch-like activation of cofilin at meiosis resumption provides a means for maintaining the stationary length of microvilli, through pointed end depolymerization of the core actin filaments, thereby replenishing the cytoplasmic monomer pool. In this model (Figure 7B), the failure to activate cofilin leads to sequestration of G-actin into microvilli, acting as a sink for monomers, at the expense of the cytoplasmic F-actin mesh which fails to reassemble. As a result, the spindle cannot migrate toward the cortex to achieve asymmetric division. Later during maturation, once the F-actin mesh has been reestablished, cofilin appears to act mainly as a regulator of actin network dynamics, with no impact on microvilli length, unless the G-actin pool is artificially raised via the disassembly of a competing actin network.

Another intriguing finding of this study is that prophase oocytes retain normal cytoplasmic F-actin dynamics despite cofilin inactivation. To explain this discrepancy, we suggest that cytoplasmic F-actin turnover during prophase arrest may rely instead on depolymerization from the barbed end. Indeed, it was proposed that at one stage of their life cycle, cytoplasmic actin filaments release Fmn2, which may leave the barbed ends unprotected from depolymerization (Montaville et al., 2014). The same scenario could explain the collapse of the cytoplamic F-actin mesh at NEBD, as it coincides with the regulated destruction of Fmn2 by the proteasome (Azoury et al., 2011). In support of this hypothesis, a truncated Cappuccino (the closest drosophila homologue to Fmn2) was shown to stabilize the fly oocyte actin mesh against disassembly by Latrunculin A, which implies that depolymerization occured at the barbed ends (Dahlgaard et al., 2007). Other actin-binding proteins may further accelerate barbed end depolymerization, such as profilin and the cofilin-related depolymerizing factor twinfilin (Montaville et al., 2014; Pernier et al., 2016; Hilton et al., 2018; Shekhar et al., 2019). Further investigations are required to verify this hypothesis and establish the molecular basis for the differential regulation of cytoplasmic F-actin during oocyte meiosis.

Notwithstanding subtle changes in morphology and density, microvilli decorate the oocyte surface across all stages of maturation, suggesting that core microvillar components are little affected by the cell cycle (Longo and Chen, 1985; Benammar et al., 2017). Yet, by interfering with cofilin activation, we uncovered a pathway for microvilli elongation relying on CDK1 activity, suggesting that microvillar actin dynamics are altered during M-phase. While the relevant CDK1 substrates remain to be characterized, a number of actin regulatory proteins have been identified as putative CDK1 targets in mitotic cells, including various formins and Cordon bleu, a candidate actin nucleator in intestinal microvilli (Dephoure et al., 2008; Grega-Larson et al., 2015). Likewise, CDK1 was shown to promote actin cable formation by the yeast formin Bni1, and Xenopus mDia2 (Miao et al., 2013). Changes in microvilli properties during oocyte maturation (e.g. accelerated treadmilling) may be a prerequisite for achieving microvilli breakdown in the polarized cortex (Dehapiot and Halet, 2013), and/or for productive interaction with the fertilizing sperm (Runge et al., 2007). In this regard, oocyte-derived exosomes, that are assumed to be of microvillar origin, are found in the perivitelline space of MII but not GV oocytes, consistent with a change in microvillar dynamics after meiosis resumption (Miyado et al., 2008; Benammar et al., 2017).

Despite a substantial activation of their cofilin pool, MII oocytes do not experience catastrophic disassembly of their actin networks, pointing to additional mechanisms to fine-tune cofilin activity. Two candidates are AIP1 and coronin, which synergize with cofilin to accelerate filament severing and disassembly, while inhibiting elongation of the severed fragments (Gressin et al., 2015; Jansen et al., 2015). While AIP1 was identified in mouse oocytes, it appears dispensable for normal oocyte function (Xiao et al., 2017; Jin et al., 2020). Srv2/CAP is another highly conserved regulator of actin turnover, acting synergistically with cofilin (Kotila et al., 2019; Shekhar et al., 2019). Accordingly, CAP1 knockdown in mouse oocytes was associated with defects in cytoplasmic F-actin, spindle migration and cytokinesis, however no microvillar outgrowth nor actin cap expansion were reported (Jin et al., 2018). Thus, the multiple actin networks that populate the oocyte may be differentially regulated by cofilin, through the assembly of distinct multi-component machineries. Additional studies are needed to characterize further these accessory proteins, as well as alternative depolymerizing factors (e.g. twinfilin), and their regulation accross the meiotic cell cycle. While oocytes provide an attractive model for microvilli plasticity and the interplay between actin dynamics and the cell cycle, it will be meaningful to investigate whether the phospho-regulation of cofilin regulates microvilli length in other cells, such as differentiating epithelial cells (Faust et al., 2019) and cells that utilize microvilli as a membrane reservoir (Figard et al., 2016).

## Materials and methods

### Mice

All animal procedures were conducted in accordance with the European directive for the use and care of laboratory animals (2010/63/EU), and approved by the local animal ethics committee under the French Ministry of Higher Education, Research and Innovation (Project licence APAFIS#11761). Mice of the MF1 strain were initially purchased from Envigo (Gannat, France) and maintained as a colony in the local animal facility. To minimize the number of animals, female mice (6-8 week old) were primed by intraperitoneal injection of 5-7 units of PMSG (Chronogest, MSD) 48h prior to GV oocyte collection. To obtain MII oocytes, mice were injected with 5 IU hCG (Chorulon, MSD) 48h after the PMSG injection.

### Oocyte collection and culture

For GV-stage oocytes, cumulus-oocyte complexes were recovered by puncturing ovarian follicles with a needle and transferred into M2 medium supplemented with 2 µM milrinone (Sigma) to maintain prophase arrest. Fully-grown GV-intact oocytes were denuded of surrounding cumulus cells by mouth pipetting. To obtain oocytes at various stages of maturation, GV oocytes were washed twice in milrinone-free M16 medium to trigger spontaneous meiosis resumption, and cultured in M16, in an atmosphere of 5% CO2 in air at 37°C, until the desired stage. Metaphase-II oocytes were recovered from the oviduct in M2 medium supplemented with 3mg/ml hyaluronidase (Sigma). The following small molecule inhibitors were used at the indicated concentration: nocodazole (5 µM; Merck Millipore), CK-666 (100 µM; Sigma), okadaic acid (2.5 µM; Enzo), cytochalasin D (100 nM; Tocris), Latrunculin A (0.3 µM; Enzo), SMIFH2 (30 µM; Sigma), flavopiridol (5 µM; Selleckchem). An equivalent amount of DMSO was used for controls. Note that in experiments involving SMIFH2, PVA (0.05%) was substituted for BSA in the culture medium.

### Cloning and mutagenesis of mouse LIMK1

Total RNA was extracted from 112 GV oocytes using Nucleospin RNA XS Kit (Macherey-Nagel), and first strand cDNA were generated with SuperScript III Reverse Transcriptase (Thermo Fisher Scientific #18080-044) and RNaseOUT (Life Technologies), using Oligo(dT)15 primers (Promega #C110A). The following primers were used to amplify the full length of LIMK1 cDNA by PCR: Forward 5’-ATGAGGTTGACGCTACTTTGT-3’, Reverse 5’-TCAGTCAGGGACCTCGGGGTG-3’. The LIMK1 fragment was next cloned into pcDNA3.1+ and the construct was validated by sequencing. Catalytically-inactive LIMK1-D460A, in which Asp-460 is replaced by Ala (Yang et al., 1998), was generated by site-directed mutagenesis using the QuickChange II Kit (Stratagene), and verified by sequencing.

### Plasmids, cRNA preparation and microinjection

Constitutively active Xenopus ADF/Cofilin XAC(A3) in pSP36T was obtained from Hiroshi Abe (Chiba University, Chiba, Japan) and subcloned into pcDNA3.1(+) for cRNA synthesis. EGFP-UtrCH in pCS2+ was a gift from William Bement (Addgene plasmid #26737; Burkel et al., 2007). Fmn2-mCherry in pCS2 was obtained from Jan Ellenberg (Euroscarf plasmid #P30607; Schuh and Ellenberg, 2008); mCherry was exchanged with EGFP in order to generate pCS2-Fmn2-EGFP. Polyadenylated cRNAs were synthesized in vitro from linearized plasmids, using the mMessage mMachine T7 or SP6 kit and Poly(A) Tailing kit (Ambion), purified with RNeasy purification kit (Qiagen), and stored at -80°C. Oocytes were injected with ∼5pl cRNA using the electrical-assisted microinjection technique, which allows for a high rate of oocyte survival (Fitzharris et al., 2018). For co-expression of LIMK1 with XAC(A3), the cRNA encoding XAC(A3) was co-injected, as a 1:1 (vol.) mix, with a 2x-concentrated LIMK1 cRNA, in order to reach LIMK1 expression level similar to oocytes injected with LIMK1 only.

### RNAi knockdown of cofilin1 transcripts

Transcripts encoding cofilin1 (*Cfl1*), the most abundant cofilin isoform expressed in oocytes (Pfender et al., 2015), were knocked down via cytoplasmic siRNA injection. Denuded prophase oocytes were microinjected with 5-10 pl of ON-TARGETplus mouse cofilin1 SMARTpool siRNA (Dharmacon #L-058638-01-0005, Horizon Discovery) diluted in RNAse-free water to a stock concentration of 100 µM. After in vitro culture in M16 medium for 22 hours, during which oocytes were maintained arrested in prophase with milrinone, total RNA was isolated from 15 oocytes using the NucleoSpin RNA XS kit (Macherey-Nagel) and processed for reverse transcription using oligo(dT)15 primers (Promega #C110A) and SuperScript III reverse transcriptase (Thermo Fisher Scientific #18080-044). The extent of cofilin1 transcript knockdown was assessed by quantitative real-time PCR performed on a QuantStudio 7 Flex system (Thermo Fisher Scientific #4485701), using the Power SYBR Green PCR master mix (Thermo Fisher Scientific #4367659). Detection of beta actin (*Actb*) transcripts was used as an internal control for normalization. Oligo pairs were as follow : cofilin1 forward : 5’- CAGACAAGGACTGCCGCTAT-3’ ; cofilin1 reverse : 5’-TTGCTCTTGAGGGGTGCATT-3’ ; beta actin forward : 5’- AGCGGTTCCGATGCCC-3’ ; beta actin reverse : 5’-CTTTACGGATGTCAACGTCACAC-3’.

### Western blot analysis

To detect endogenous cofilin and phospho-cofilin, 50-100 oocytes were harvested at the indicated stage of maturation, washed in PBS supplemented with 1% polyvinylpyrrolidone (Sigma) and lysed in Laemmli 2× Concentrate Sample Buffer (Sigma). Lysates were immediately frozen at -80°C until use. After thawing and denaturation at 95°C for 5 min, proteins were separated by SDS-PAGE on NuPAGE Novex 4-12% Bis-Tris gels (Life Technologies) and transferred onto PVDF membranes (Amersham) in Tris-glycine-10% ethanol. Membranes were blocked with PBST (PBS containing 0.1% Tween 20) supplemented with 5% BSA or 5% non-fat dry milk for 3 h at room temperature, and then incubated with primary antibodies at 4°C overnight. After washing with PBST three times, membranes were incubated with secondary antibodies conjugated to horseradish peroxidase for 1 h at room temperature, followed by wash with PBST three times. Blots were developed using SuperSignal West DURA Extended Duration Substrate (Thermo Fisher Scientific) following the manufacturer’s instructions. Semi-quantitative densitometric analysis of band intensities was performed using the Gels command in FIJI open source software. The following primary antibodies were used: rabbit anti-cofilin (Cytoskeleton ACFL02; 1:500), rabbit anti-Ser3-phosphorylated cofilin (Cell Signaling Technology #3311; 1:500), and a mouse anti-GAPDH (Abcam ab8245; 1:5000), to use as a control for protein loading. Secondary antibodies were HRP-conjugated goat anti-rabbit IgG (G-21234; 1:25000; Life Technologies) and goat anti-mouse IgG (G-21040; 1:25000; Life Technologies).

### Immunofluorescence

Oocytes were fixed for 16 min at room temperature with paraformaldehyde 3% in PBS, freshly prepared from a 16% methanol-free paraformaldehyde solution (Electron Microscopy Sciences). Fixed oocytes were permeabilized with 0.25% Triton X-100 (Sigma) in PBS for 15 min, blocked with 3% BSA (Sigma) in PBS for 3 h, and incubated overnight at 4°C with primary antibodies diluted in PBS-BSA 3%. On the next day, oocytes were washed in PBS-BSA 1% and incubated with secondary antibodies diluted in PBS-BSA 1%, for 45 min at 37°C. The following primary antibodies were used at the indicated dilution: anti-tubulin (rat monoclonal, Abcam #ab6161; 1:200), anti-phospho-ERM (rabbit monoclonal, Cell Signalling Technology #3726; 1:100), anti-CD9 (rat monoclonal, Santa Cruz sc-18869; 1:100), anti-DAAM1 (goat polyclonal, Santa Cruz sc-55929; 1:100). Secondary antibodies were Alexa Fluor 488-conjugated donkey anti-goat, donkey anti-rabbit and goat anti-rat (all 1:1000; Invitrogen). Chromatin was stained with To-Pro-3 (Invitrogen T3605).

### Chromosome spreading

To assess homologous segregation, LIMK1-expressing oocytes that had been cultured overnight in M16 medium were processed for chromosome spreading based on a previously published protocol (Roberts et al., 2005). Briefly, oocytes were treated with a hypotonic solution (fresh 1% sodium citrate) for 5 min, then individually transferred into small (5 µl) drops of fixative (5:1:4 of methanol:acetic acid:water) on a glass-bottom dish (Mattek). After the zona pellucida had gradually dissolved, 10 µl of fixative (3:1 of methanol:acetic acid) were gently dropped onto the oocytes (repeated 2-3 times), followed by air-drying at room temperature. Chromosomes were labeled with Hoechst 33342 (10 µg/ml in water;) for 30 min followed by wash with water. Images of chromosome spreads were acquired using an inverted Leica DMI4000B microscope equipped with a 365nm LED module, using a 63x oil-immersion objective.

### DNAse I staining for detection of cytoplasmic G-actin

To assess the levels of G-actin in the cytoplasm, oocytes were stained with DNAse I according to a previously published protocol (Yu et al., 2014), with minor modification. Briefly, oocytes were fixed in 4% paraformaldehyde in fixation buffer (130 mM KCl, 3 mM MgCl2, 25 mM HEPES pH 7, 0.15% glutaraldehyde), for 30 min at room temperature. Fixed oocytes were permeabilized with 0.5% Triton X-100 for 15 min and blocked with 3% BSA in PBS for 2 h. Oocytes were next incubated with Alexa Fluor 488-conjugated DNAse I (Thermo Fisher Scientific #D12371) diluted to 10 µg/ml in PBS-BSA 3%, for 30 min at room temperature, followed by wash in PBS-BSA 3%. Confocal images were acquired (see below) and DNAse 1 fluorescence was measured in a 10×10 µm cytoplasmic region, away from the GV or spindle area. To correct for background fluorescence, we performed triton-extraction on live oocytes before fixation (Kan et al., 2011), in order to remove all soluble G-actin. Thus, live oocytes were incubated for 10 min at room temperature in extraction buffer (100 mM KCl, 20 mM MgCl2, 3 mM EGTA, 20 mM HEPES pH 6.8, 0.1% Triton X-100), washed in PBS, and then processed for fixation as described above. At least five oocytes were extracted in each experimental series, in order to calculate an averaged background fluorescence value, that was subtracted from raw values in non-extracted oocytes.

### F-actin staining and quantification in fixed oocytes

F-actin was stained with Alexa Fluor 568-phalloidin (Life Technologies). The density of the cytoplasmic F-actin network in MI (NEBD + 6h) oocytes was evaluated by measuring the mean fluorescence intensity in a 10×10 µm region of interest using the Leica LAS AF Lite 2.6.0 software. Care was taken to avoid the spindle region and the elongated cytoplasmic actin rootlets. To calculate the density of elongated microvilli in LIMK1-expressing MI oocytes, a single confocal optical section (thickness 1 µm) obtained at the equator of the oocyte, was analyzed. Elongated microvillar actin bundles were counted across the oocyte perimeter, and their density was expressed as per 10-µm perimeter. Only elongated structures showing bidirectional elongation (outward microvilli + inward rootlet) were considered for the analysis. In fixed MII oocytes, the thickness of the actin cap (defined as the distance spanning from the submembrane cortical actin to the surface of the spindle), and of the cortical actin layer in the opposite cortex, were measured offline using Leica LAS AF Lite 2.6.0.

### Analysis of microvilli length with FiloQuant

We used the imageJ plugin FiloQuant (Jacquemet et al., 2017) to detect elongated microvilli and measure their length. A single confocal section (thickness 1 µm) obtained at the equator of the oocyte, was processed using the single image analysis mode, and outward microvilli were resolved as illustrated in Figure S3. First, an unsharp filter was applied in FIJI to facilitate microvilli detection. Next, the oocyte contour was manually outlined, and the cytoplasmic region was filled using the Fill function, in order to allow for thresholding of the cell edge. Transzonal projections contaminating the F-actin signal were manually outlined and removed using the Clear function. The resulting image was processed with Filoquant in the single image mode. Threshold parameters for cell edge and microvilli detection were adjusted iteratively for best results. A lower cut-off at 1-µm was applied to minimize detection of irrelevant bright spots originating mostly from numerous microvilli fragments crossing the plane of the confocal frame (“broken” microvilli).

### F-actin labeling and quantification in live oocytes

F-actin networks were labeled using EGFP-UtrCH expressed via cRNA microinjection. For GV and MI oocytes, cRNA injection was performed in GV oocytes arrested in prophase with milrinone, followed by culture for 3 hours in M16-milrinone, to allow for protein expression. For MII oocytes, cRNA injection was followed by a 3-hour culture in M16 for protein expression. In LIMK1-expressing oocytes, the cRNA encoding LIMK1 was co-injected, as a 1:1 (vol.) mix, with a 2x-concentrated EGFP-UtrCH cRNA, in order to reach a similar expression level of the F-actin probe in LIMK1-injected oocytes and control oocytes. In some experiments, chromosomes were labeled with SiR-DNA (Spirochrome), added to the culture medium to a final concentration of 0.5 µM. Spatiotemporal image correlation spectroscopy (STICS) analysis was performed in FIJI, using the dedicated plugin written by Jay Unruh (Stowers Institute for Medical Research, Kansas City, MO).

### Confocal imaging, image processing and analysis

Oocytes were placed on glass-bottom dishes (MatTek, Ashland, MA) and imaged with a Leica SP5 or SP8 confocal microscope, using a 63x oil-immersion objective. The thickness of optical sections was set at 1 µm. For live imaging, temperature was maintained at 37°C using a stage top incubator (model INUBG2E-GSI, Tokai Hit, Shizuoka-ken, Japan) fitted on the microscope stage. Confocal settings were kept identical in control and experimental (e.g. LIMK1 overexpression, drug treatment) conditions. Confocal images and time-lapse movies were processed with FIJI. Image sharpness was enhanced by applying an unsharp mask filter in FIJI. Kymographs were generated with the Multi Kymograph tool in FIJI. Fluorescence intensities (in arbitrary units) were expressed as mean +/- SD.

### Statistical analysis

All experiments were repeated independently at least three times. Mean, standard deviation (SD) or standard error (SE), and statistical significance (P<0.05) were calculated using the Student’s t test or Mann-Whitney test in Origin (OriginLab). Line graphs and bar graphs were generated in Excel or Origin. Box plots were generated in Origin, and show the mean (square), median (line), 25^th^, 75^th^ percentiles (box) and 5^th^, 95^th^ percentiles (whiskers).

## Acknowledgements

We thank Prof. Hiroshi Abe for the XAC(A3) plasmid and Jay Unruh for advice with STICS analysis. We are grateful to the staff of the ARCHE-Biosit animal facility and MRIC-Biosit microscopy facility for technical assistance and expert advice.

## Author contributions

GH conceived and supervised the study. AB, BD and GH designed and performed experiments, and analysed data. GH prepared the figures and wrote the manuscript with input from AB and BD.

---

**Figure S1.**
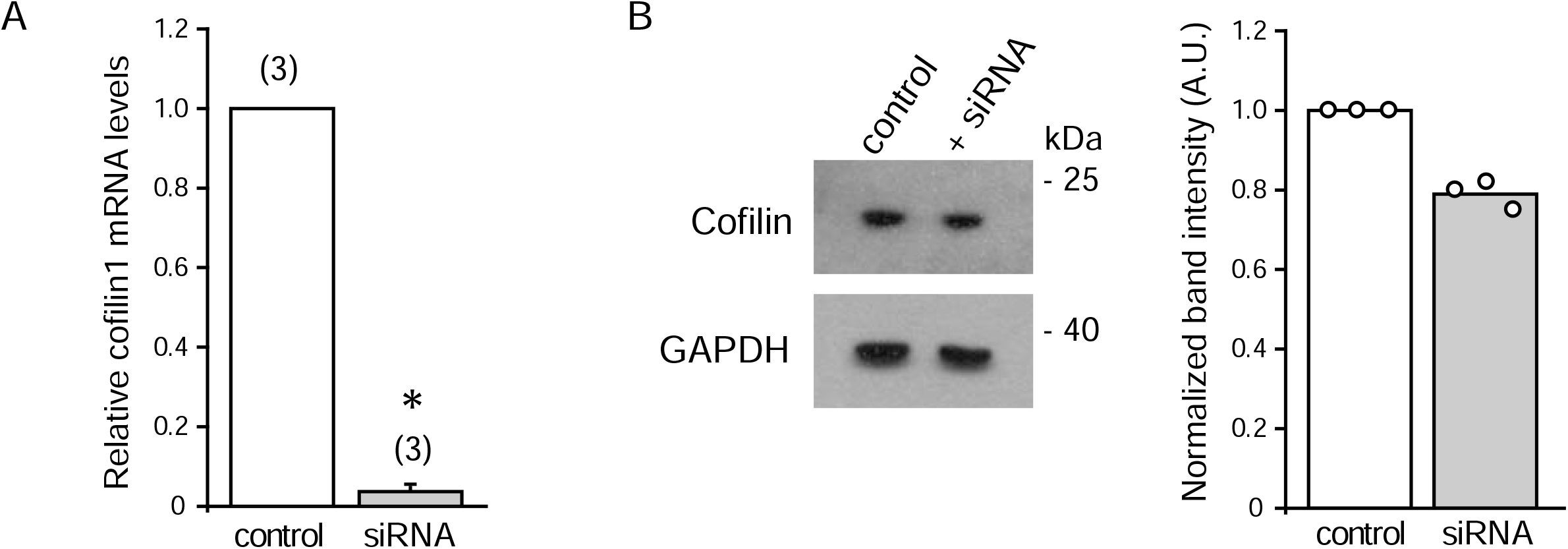
Depletion of cofilin mRNA by RNA interference. GV oocytes were injected with siRNA against cofilin1 and cultured for 22 h with milrinone to maintain prophase arrest. (A) Quantification of cofilin1 mRNA levels by quantitative real time PCR, using beta actin mRNA as an internal control for normalization. Data are means +/- SD of three independent experiments. * : P<10^−5^. (B) Detection of total cofilin levels in GV oocyte lysates, by western blot. GAPDH detection was used as a loading control. The bar graph shows the quantification of band intensities after normalization to the GAPDH signal. The experiment was repeated three times.

**Figure S2.**
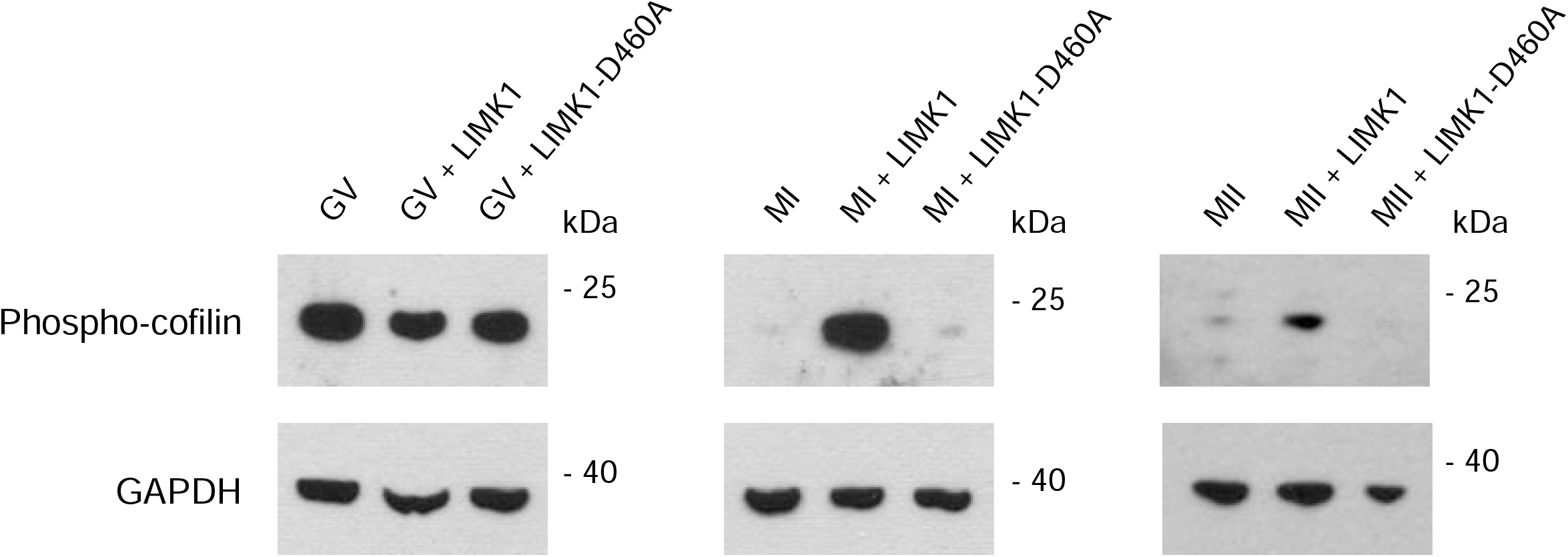
LIMK1 overexpression increases phospho-cofilin levels. Phospho-cofilin was detected in lysates from 70-80 GV (left panel), MI (middle panel) and MII (right panel) oocytes. Overexpression of LIMK1 (+LIMK1), or catalytically-inactive LIMK1-D460A (+LIMK1-D460A) was achieved by microinjection of the corresponding cRNA. For GV oocytes (left panel), injection was performed at the GV stage and oocytes were cultured in vitro for 3 h before lysis. For MI oocytes, injection was performed at the GV stage and oocytes were cultured in vitro until the MI (NEBD+6h) stage. For MII oocytes, injection was performed in freshly isolated MII oocytes, which were cultured in vitro for 3 h before lysis. Note that MI oocytes expressed LIMK1 for an extended period of time (7-8 h), which may explain the higher band intensity in comparison with MII oocytes. Detection of GAPDH was used as a loading control. Data are representative of at least 3 similar observations for LIMK1-overexpression and two similar observations for LIMK1-D460A overexpression.

**Figure S3.**
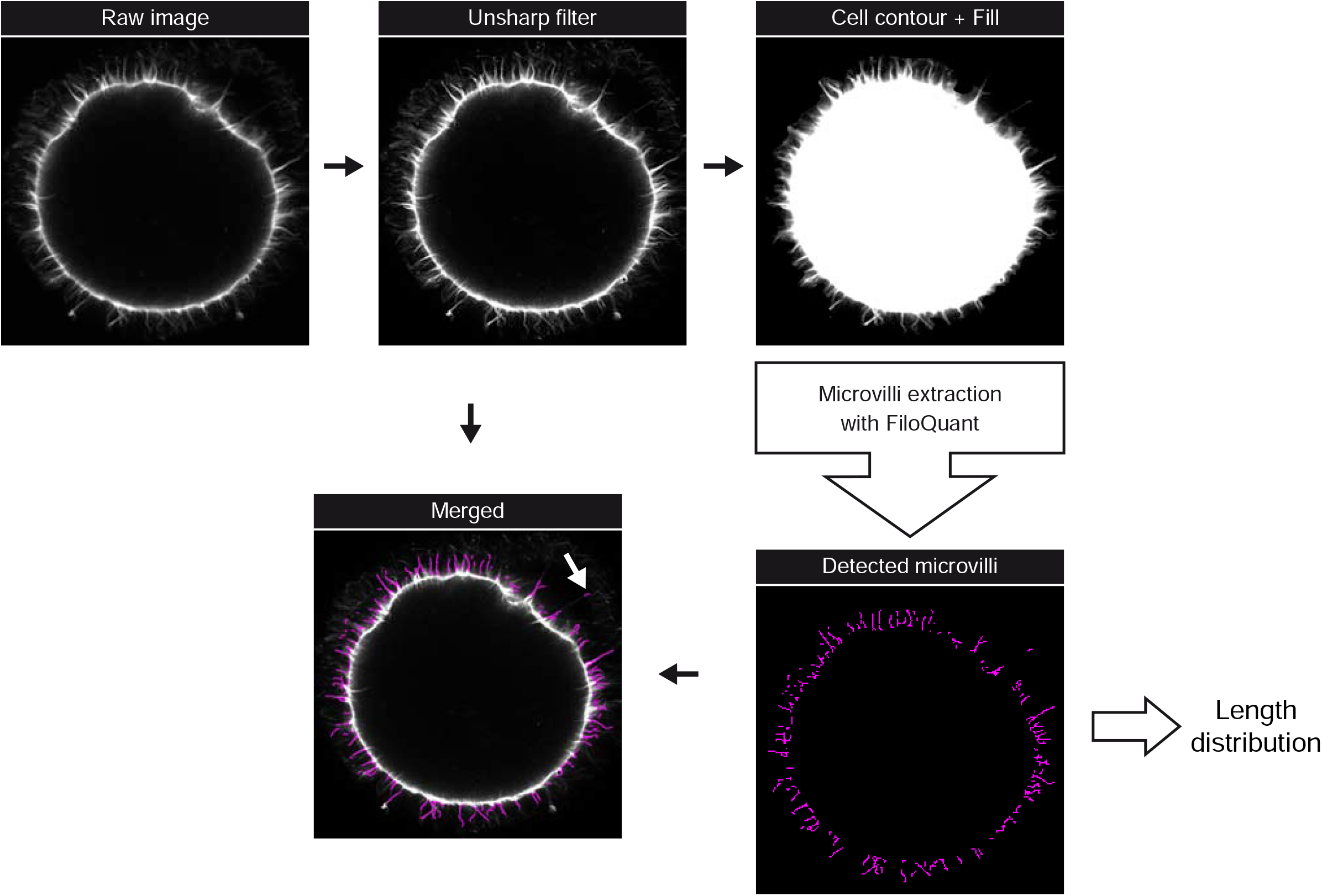
Detection of elongated microvilli with Filoquant. Filoquant (Jacquemet et al., 2017) was used to detect microvilli in fixed oocytes stained with Alexa Fluor 568-phalloidin. The image processing workflow is shown for a LIMK1-expressing MI oocyte treated with CK-666. Raw confocal images (thickness 1 µm) were first processed with Fiji to sharpen the fluorescence signal and delineate the cell contour. Microvilli (magenta) were resolved with Filoquant in the single image analysis mode, by iteratively adjusting the settings for cell edge detection and “filopodia” detection. In the merged image, the white arrow points to a “broken” microvillus fragment.

**Figure S4.**
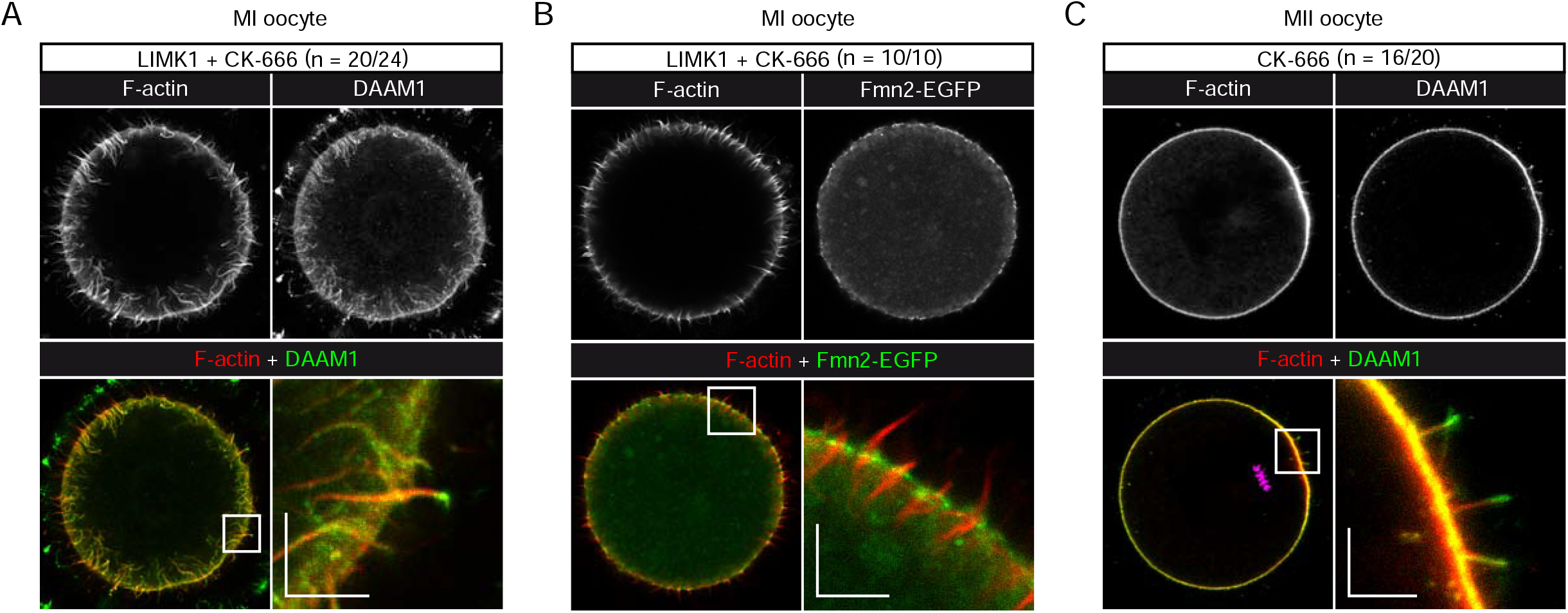
DAAM1 decorates elongated microvilli and accumulates at the tips. (A,B) LIMK1-expressing MI oocytes treated with CK-666 (100 µM) and fixed at MI (NEBD+6h). (C) MII oocyte treated with CK-666 (100 µM) for 3h before fixation. In (A) and (C), oocytes were immuno-stained for DAAM1 (green). In (B), oocytes were coinjected with cRNA encoding Fmn2-EGFP (green). F-actin was stained with Alexa Fluor 568-phalloidin (red). Chromosomes in (C) are labeled with TO-PRO-3 (magenta). Expanded views of cortical areas with elongated microvilli (white boxes) are shown. The number of oocytes examined showing a similar pattern is indicated in parentheses. Scale bars represent 5 µm.

## Supplementary movie legends

**Movie 1. Cofilin is dispensable for cytoplasmic F-actin dynamics in GV oocytes**. Time-lapse confocal imaging (one image every 30 sec) of F-actin networks in GV oocytes expressing EGFP-UtrCH together with catalytically-inactive LIMK1^D460A^ (Control, left), or LIMK1 (right). Spontaneous meiosis resumption was triggered by washing off milrinone. Chromosomes are labeled with Sir-DNA (magenta). Confocal section thickness is 1 µm.

**Movie 2. Cytoplasmic F-actin is absent in LIMK1-expressing MI oocytes**. Time-lapse confocal imaging (one image every 30 sec) of F-actin networks in oocytes expressing EGFP-UtrCH. Left to right: uninjected MI oocyte (Control), LIMK1-expressing MI and LIMK1^D460A^-expressing MI oocytes. Chromosomes are labeled with Sir-DNA (magenta). Confocal section thickness is 1 µm.

**Movie 3. Arp2/3 inhibition exacerbates microvillar actin elongation**. Confocal z-stack of a fixed LIMK1-expressing MI oocyte labeled for F-actin with Alexa Fluor 568-phalloidin. CK-666 (100 µM) was added to the culture medium at the onset of meiosis resumption (milrinone wash). Chromosomes are labeled with TO-PRO-3 (magenta). Confocal section thickness is 1 µm. The outermost layer shows a weak labeling of transzonal projections crossing the zona pellucida.

**Movie 4. Elongated microvilli are enriched in P-ERM**. Confocal z-stack of a LIMK1-expressing MI oocyte treated with CK-666 (100 µM) and labeled for F-actin (Alexa Fluor 568-phalloidin, red) and P-ERM (green). Chromosomes are labeled with TO-PRO-3 (magenta). Confocal section thickness is 1 µm.

**Movie 5. F-actin dynamics in MII and LIMK1-expressing MII oocytes**. Time-lapse confocal imaging (one image every 30 sec) of F-actin networks in a control MII oocyte (left) and a LIMK1-injected MII oocyte (right) expressing EGFP-UtrCH to label actin filaments. Confocal section thickness is 1 µm.

**Movie 6. Ectopic microvilli are enriched in CD9**. Confocal z-stack of an MII oocyte treated with CK-666 (100 µM) for 3h, then fixed and labeled for F-actin (Alexa Fluor 568-phalloidin, red) and CD9 (green). Chromosomes are labeled with TO-PRO-3 (magenta). The red halo at the start of the series is due to the F-actin signal in the first polar body. Confocal section thickness is 1 µm.

